# The *Drosophila* dystonia gene homolog *Neurocalcin* facilitates sleep by inhibiting a movement-promoting neural network

**DOI:** 10.1101/159772

**Authors:** Ko-Fan Chen, Angélique Lamaze, Patrick Krätschmer, James E.C. Jepson

## Abstract

Primary dystonia is a hyperkinetic movement disorder linked to altered dopaminergic signaling and synaptic plasticity in regions of the brain involved in motor control. Mutations in *HPCA*, encoding the neuronal calcium sensor Hippocalcin, are associated with primary dystonia, suggesting a function for Hippocalcin in regulating the initiation and/or maintenance of activity. However, such a role for Hippocalcin or Hippocalcin homologs has yet to be demonstrated in vivo. Here we investigate the cellular and organismal functions of the *Drosophila* Hippocalcin homolog Neurocalcin (NCA), and define a role for NCA in promoting sleep by suppressing nighttime hyperactivity. We show that NCA acts in a common pathway with the D1-type Dop1R1 dopamine receptor and facilitates sleep by inhibiting neurotransmitter release from a multi-component activity-promoting circuit. Our results suggest conserved roles for Hippocalcin homologs in modulating motor control through dopaminergic pathways, suppressing aberrant movements in humans and inappropriate nighttime locomotion in *Drosophila*.

## Introduction

Primary dystonia is characterized by repetitive involuntary movements or sustained abnormal postures, and represents the third most common movement disorder after benign tremor and Parkinson’s disease (Fahn, 1988; Wenning et al., 2005). Dystonia has been linked to altered neurotransmission and/or plasticity of circuits within the basal ganglia, a group of nuclei involved in action initiation and maintenance, as well as associated inputs/outputs (Calabresi et al., 2016; Karimi and Perlmutter, 2015; Pappas et al., 2015; Weisheit and Dauer, 2015). However, the molecular pathways underlying this disorder remain unclear.

Identifying genetic loci linked to hereditary forms of primary dystonia represents a useful strategy to uncover such pathways. Mutations in *GNAL*, encoding Gα_olf_, have been linked to primary dystonia (Fuchs et al., 2013), and since the Gα_olf_ G-protein α-subunit couples to D1-type dopamine receptors, this finding supports a link between dystonia and altered dopaminergic signaling (Karimi and Perlmutter, 2015). Indeed, current models suggest that the basal ganglia regulate motor control through two distinct circuits, the direct and indirect pathways, that are defined by expression of D1- and D2-type type dopamine receptors respectively (Gerfen and Surmeier, 2011; Tecuapetla et al., 2016). These pathways antagonistically influence activity of large-scale brain networks (Lee et al., 2016), and an imbalance of direct versus indirect pathway signaling has been proposed to cause dystonia (Breakefield et al., 2008; Peterson et al., 2010; Tanabe et al., 2009; Yokoi et al., 2015). Mutations in *dTorsin*, the *Drosophila* homolog of the dystonia gene *TOR1A*, reduce dopamine levels by suppressing expression of the dopamine synthesis factor GTP cyclohydrolase (Wakabayashi-Ito et al., 2015; Wakabayashi-Ito et al., 2011), further linking dystonia to altered dopamine signaling.

Recent studies have identified an array of further loci linked to primary dystonia, including *THAP1*, *ANO3*, *KCTD17* and *HPCA* (Charlesworth et al., 2015; Charlesworth et al., 2012; Fuchs et al., 2009; Mencacci et al., 2015). However, in vivo data linking these genes to motor control and/or dopaminergic pathways is limited. Interestingly, mutations in the *Drosophila KCTD17* homologue *insomniac* result in reduced sleep (i.e increased movement), a phenotype that can be rescued by inhibiting dopamine synthesis (Pfeiffenberger and Allada, 2012; Stavropoulos and Young, 2011). Thus, we sought to examine whether other dystonia gene homologs in *Drosophila* also impacted sleep, beginning with the *Drosophila HPCA* homolog *Neurocalcin* (*Nca*).

NCA and Hippocalcin (the *HPCA* gene product) act as neuronal calcium sensors, cytoplasmic proteins that bind to calcium via EF hand domains and translocate to lipid membranes through a calcium-dependent myristoylation switch (Burgoyne and Haynes, 2012). This switch modulates interactions with membrane-bound ion channels and receptors, altering target function or localization (Braunewell and Klein-Szanto, 2009; Burgoyne and Haynes, 2012). Missense mutations in *HPCA* have been linked to DYT2 primary isolated dystonia, which predominantly affects the upper limbs, cervical and cranial regions (Charlesworth et al., 2015). The index N75K mutation in *HPCA* alters a neutral asparagine residue in the second EF hand to a positively charged lysine, likely interfering with calcium binding in a loss-of-function manner.

Hippocalcin undertakes pleiotropic roles in mammalian neurons (Braunewell and Klein-Szanto, 2009), including acting as a calcium sensor to gate the slow afterhyperpolarisation (sAHP), a calcium-dependent potassium current, and facilitating NMDA receptor endocytosis during LTD (Jo et al., 2010; Tzingounis et al., 2007). In *Drosophila*, NCA has previously been shown to be broadly expressed throughout the adult nervous system (Teng et al., 1994), yet the neuronal and organismal roles of NCA are unknown. Here we show that NCA is required to promote sleep by suppressing locomotor activity during the night. We demonstrate that NCA acts in a common pathway with the D1-type Dop1R1 dopamine receptor and identify a multi-component wake-promoting neuronal network in which NCA facilitates sleep by suppressing synaptic release. Thus, we propose that Hippocalcin homologs play conserved roles in regulating aspects of motor control.

## Results

### Generation of *Nca* knockout flies

The *Drosophila* genome contains a single *HPCA* homolog, *Neurocalcin* (*Nca*). Hippocalcin and NCA share > 90% amino-acid identity (Figure 1 – figure supplement 1A), suggesting conservation of function. To investigate the neuronal and behavioral roles of NCA we generated a *Nca* null allele by replacing the entire *Nca* locus (including 5’ and 3’ UTRs) with a mini-*white*^*+*^ marker sequence using ends-out homologous recombination (Baena-Lopez et al., 2013) (Figure 1A, Figure 1 – figure supplement 1B-C). The mini-*white*^*+*^ sequence is flanked by loxP sites, allowing removal by Cre recombinase and leaving single attP and loxP sites in place of the *Nca* locus (Figure 1A). As expected, no *Nca* mRNA expression was detected in homozygotes for the deleted *Nca* locus (Figure 1 – figure supplement 1D, E). Thus, we term this allele *Nca*^KO^ (*Nca* knockout).

**Figure 1.**
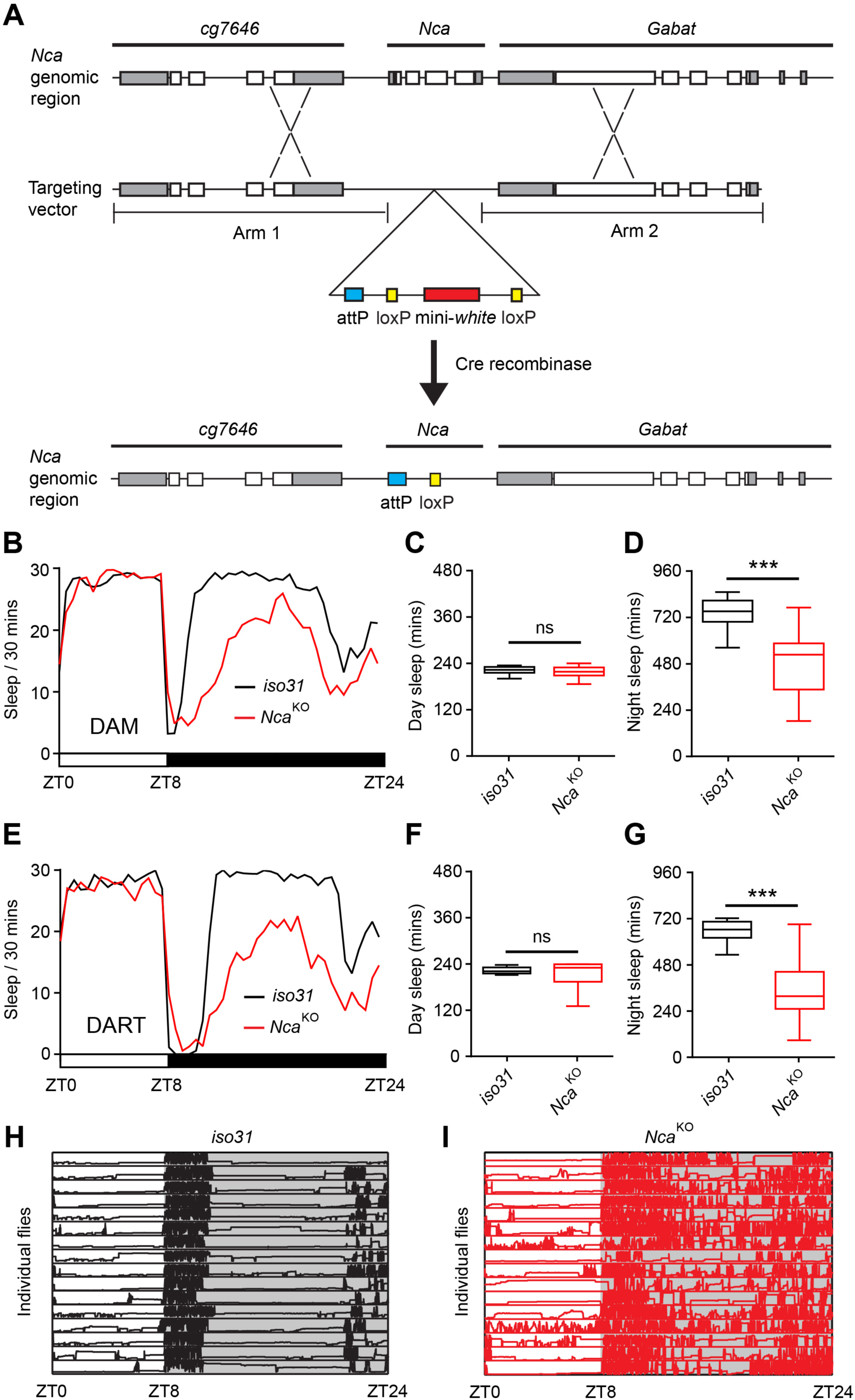
*Neurocalcin* (*Nca*) knockout flies exhibit enhanced night activity and reduced sleep. (A) Schematic illustration of the procedure used to generate a *Nca* knockout allele. Homologous arms upstream (Arm 1) and downstream (Arm 2) of the *Nca* locus are indicated. Following homologous recombination, the endogenous *Nca* locus is replaced by a cassette containing the mini-*white* selection marker (red bar), and attP (blue bar) and loxP sites (yellow bars). The mini-*white* cassette was subsequently removed via Cre-loxP recombination. (B) Mean sleep levels in 8L: 16D conditions for *Nca*^KO^ adult males and *iso31* controls measured using the *Drosophila* Activity Monitor (DAM). (C-D) Median day (C) and night (D) sleep levels in the above genotypes. (E) Mean sleep levels in 8L: 16D conditions for *Nca*^KO^ adult males and *iso31* controls measured by the video-based *Drosophila* ARousal Tracking system (DART). (F-G) Median day (F) and night (G) sleep levels in the above genotypes. Data are presented as Tukey box plots. The 25^th^, Median, and 75^th^ percentiles are shown. Whiskers represent 1.5 x the interquartile range. Identical representations are used in all subsequent box plots. B-D: n = 32 per genotype; E-G: n = 16. (H-I) The longitudinal movement for individual *iso31* (H) and *Nca*^KO^ (I) flies are shown as rows of traces plotting vertical position (Y-axis) over 24 h (X-axis) under 8L: 16D condition. ***p < 0.001, ns - p > 0.05, Mann-Whitney U-test.

### *Nca* knockout flies exhibit reduced night sleep

Following outcrossing into an isogenic *iso31* control background, *Nca*^KO^ homozygotes were viable to the adult stage, allowing us to test whether NCA impacts locomotor control. To do so, we first used a high-throughout yet low resolution system, the *Drosophila* Activity Monitor (DAM) (Pfeiffenberger et al., 2010a), which counts locomotor activity via perturbations of an infrared beam intersecting a glass tube housing individual flies. Using this system, under 12 h light: 12 h dark conditions (12L: 12D, 25°C) we found that *Nca*^KO^ males exhibited two distinct locomotor phenotypes. Firstly, reduced maximal locomotor activity in response to lights-off (also termed the startle response or masking) (Figure 1 – figure supplement 2A, B). Secondly, increased locomotor activity during the night but not the day (Figure 1 – figure supplement 2A, C, D). This later phenotype was suggestive of reduced night sleep. We therefore quantified sleep levels in *Nca*^KO^ males and controls using a 5 min period of inactivity to define a sleep bout – the standard definition in the field (Pfeiffenberger et al., 2010b). Indeed, *Nca*^KO^ males exhibited reduced night sleep but not day sleep relative to controls (Figure 1 – figure supplement 3). Interestingly, the impact of removing NCA on night sleep appeared further enhanced by shortening photoperiod, such that night (but not day) sleep was substantially reduced in *Nca*^KO^ males under 8L: 16D (Figure 1B-D). Similarly to 12L: 12D, peak locomotor activity following lights-off was reduced in *Nca*^KO^ males under 8L: 16D, and the number of beam breaks during the normally quiescent period of the night was increased (Figure 1 – figure supplement 4).

To obtain a higher resolution analysis of locomotor patterns in *Nca*^KO^ flies we next utilized a video-tracking method - the DART (*Drosophila* ARousal Tracking) system (Faville et al., 2015). Continual video-monitoring of *Nca*^KO^ males and *iso31* controls confirmed robust sleep loss in *Nca*^KO^ males specifically during the night (Figure 1E-G) accompanied by significantly shortened sleep bouts (Figure 1 – figure supplement 5). By examining locomotor patterns in individual flies, we found that *Nca*^KO^ males consistently displayed prolonged activity relative to controls following lights-off and frequent bouts of movement even in the middle of the night, a period of quiescence in *iso31* controls (Figure 1H, I). The velocity of locomotion in *Nca*^KO^ males was also significantly increased relative to moving controls in the normally quiescent period of the night (4-12 h after lights-off), whereas peak velocity during the startle response to lights-off was reduced (Figure 1 – figure supplement 6), in agreement with data collected using the DAM system. Collectively, the above data suggest that complete loss of NCA results in both locomotor deficits and profound nighttime hyperactivity, resulting in reduced peak levels of activity and night sleep.

### NCA is required in neurons to promote night sleep

To identify the cellular substrates in which NCA acts to regulate locomotion and sleep we performed cell-specific knockdown of *Nca* expression using transgenic RNA interference (RNAi). NCA has been shown to be widely expressed in neuropil regions throughout the *Drosophila* brain (Teng et al., 1994). Thus, we initially examined neurons as a potential cellular candidate. We used three independent RNAi lines (*kk108825, hmj21533* and *jf03398,* termed *kk*, *hmj* and *jf* respectively) to reduce *Nca* expression in adult male neurons using the pan-neuronal *elav*-Gal4 driver. The *kk* and *jf* dsRNAs target a partially overlapping sequence of *Nca*, whereas *hmj* targets a distinct, non-overlapping upstream sequence (Figure 2 – figure supplement 1A). For each RNAi line, we confirmed reduced *Nca* expression using qPCR (Figure 2 – figure supplement 1B). Transcription of *Nca* occurs from promoter regions shared with the downstream locus *cg7646*, yet *cg7646* transcription was not affected by *Nca* RNAi (Figure 2 – figure supplement 1C), nor were any common off-target mRNAs predicted for the *kk*, *hmj* or *jf* dsRNA hairpins (data not shown). Importantly, under 8L: 16D conditions, pan-neuronal expression of all three *Nca* RNAi lines specifically reduced night sleep in males as measured by the DAM system (Figure 2A-C, Figure 2– figure supplement 2). However, *Nca* knockdown did not result in a reduction in peak locomotor activity as observed in *Nca*^KO^ males (Figure 2 – figure supplement 3). Thus, neuronal NCA predominantly impacts sleep/activity during the night, with NCA potentially acting in other cell types to regulate peak locomotor levels. We therefore focused on elucidating the genetic pathways and neuronal circuits in which NCA acts to promote night sleep.

**Figure 2.**
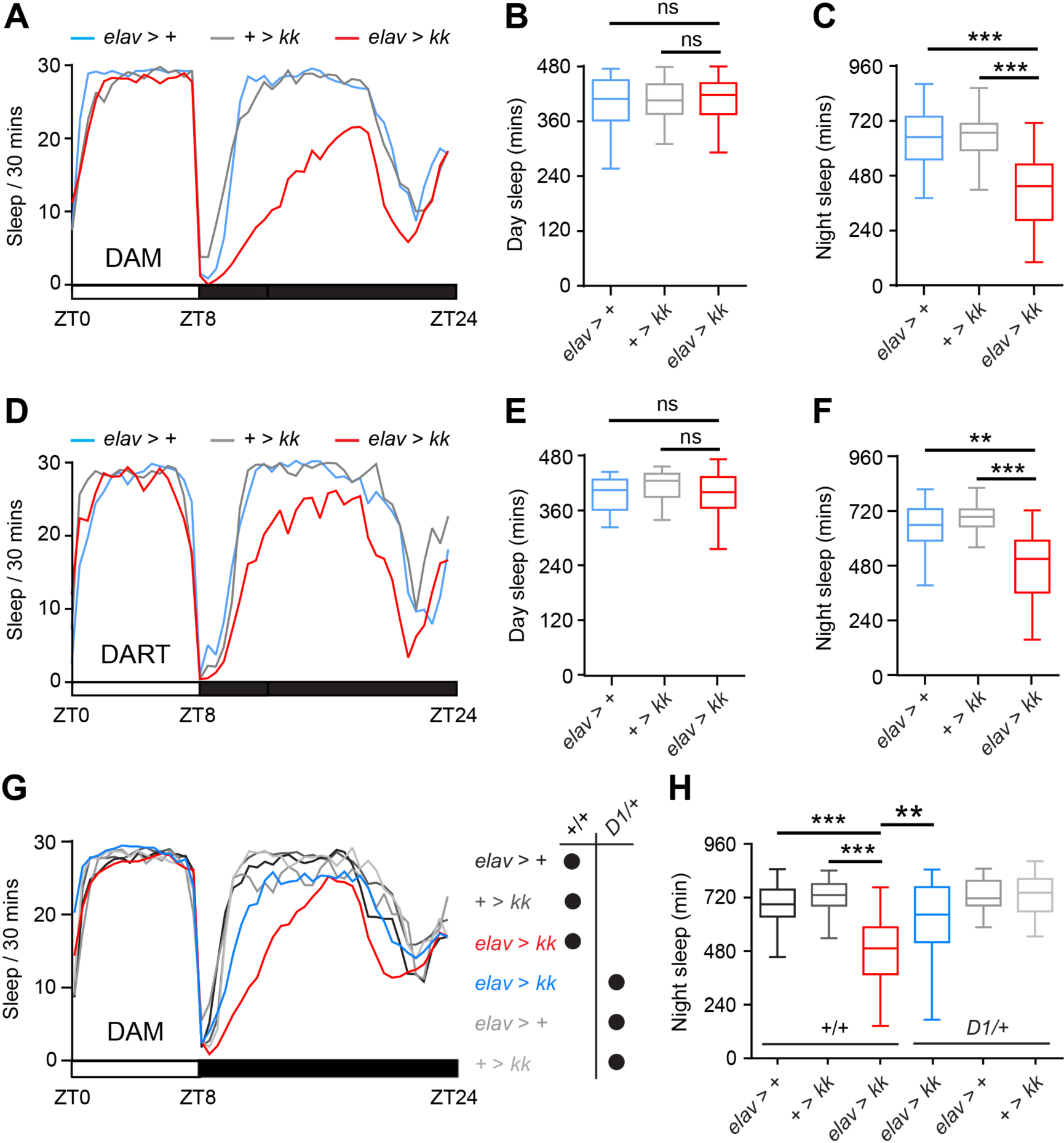
*Nca* and *Dop1R1* act in a common pathway to regulate night sleep. (A) Mean sleep levels measured using the DAM system under 8L: 16D conditions for pan-neuronal *Nca* knockdown (*Nca*^KD^) male adult flies (*elav > kk*) and associated controls (*elav*-Gal4 driver or *kk* RNAi transgene heterozygotes). (B-C) Median day and night sleep levels in the above genotypes. n = 54-55. (D) Mean sleep levels measured using DART system in 8L: 16D conditions for above genotypes. (E-F) Median day and night sleep levels in the above genotypes. n = 20 per genotype. (G-H) heterozygosity for the null or strongly hypomorphic Dop1R1 allele *Dop1R1*^MI03085-GFST.2^ (*D1*/+) suppressed sleep loss in *Nca*^KD^ adult males, but did not alter sleep in control males (p > 0.05, between +/+ and *D1*/+ backgrounds for *elav > +* or *+ > kk* flies). Mean sleep patterns in 8L: 16D conditions are shown in (G). Median total night sleep levels are shown in (H). *elav > kk, D1/+*: n = 32; *elav > +, D1/+*: n = 15; *+ > kk, D1/+*: n = 32; *elav > kk*: n = 48*; elav > +*: n = 47; *+ > kk*: n = 48. **p < 0.01, ***p < 0.001, ns – p < 0.05, Kruskal-Wallis test with Dunn’s post-hoc test.

To test whether sleep loss caused by neuronal *Nca* knockdown flies was due to an indirect effect on the circadian clock, we examined whether *Nca* knockdown altered circadian patterns of locomotor activity under constant dark conditions (Figure 2 – figure supplement 4A, B). Importantly, knockdown of *Nca* in neurons did not alter circadian rhythmicity (Figure 2 – figure supplement 4A, B), nor did *Nca* expression cycle in whole fly heads (Figure 2 – figure supplement C). Thus, it is unlikely that sleep loss caused by neuronal *Nca* knockdown is due to circadian clock dysfunction. We extended our initial findings in *Nca* knockdown males harbouring the pan-neuronal *elav*-Gal4 driver and found that night sleep loss due to neuronal *Nca* knockdown was also observed in adult virgin females (Figure 2 – figure supplement 5), and in male flies expressing the *kk Nca* RNAi using distinct pan-neuronal (*nsyb*-Gal4) or broadly expressed (*insomniac*-Gal4) drivers (Figure 3A). In contrast, expression of the *kk Nca* RNAi in muscle cells did not alter night sleep in adult males (Figure 2 – figure supplement 6), supporting the premise that NCA acts in the nervous system to regulate sleep. Video tracking further confirmed that neuronal expression of *kk Nca* RNAi reduced night sleep (Figure 2D-F). Collectively, the above data demonstrate that NCA acts in neurons to promote night sleep in *Drosophila*. For simplicity, we use the *kk Nca* RNAi for all subsequent experiments, and refer to flies expressing *kk Nca* RNAi under *elav*-Gal4 as *Nca*^KD^ (*Nca* knockdown). Given the similar results obtained using the DAM and DART systems for both *Nca*^KO^ and *Nca*^KD^ flies, we use DAM as a high-throughput method for all sleep measurements detailed below, which are performed in 8L: 16D conditions at 25°C.

**Figure 3.**
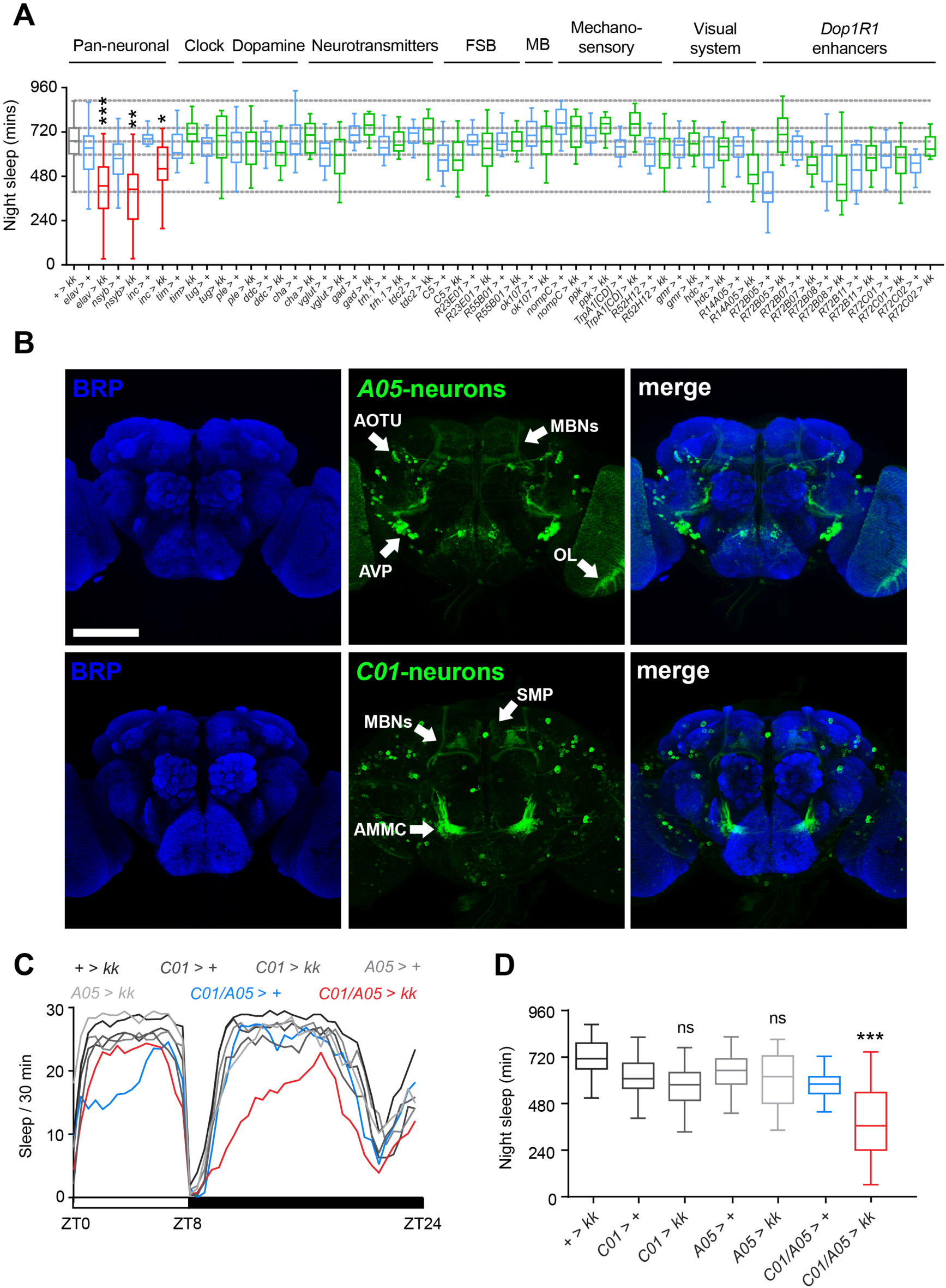
NCA acts in a two distinct neural networks to regulate night sleep. (A) Transgenic RNAi-based mini-screen to identify key NCA-expressing neurons. NCA knockdown with broadly expressed drivers resulted in reduced night sleep in adult males under 8L: 16D conditions. In contrast, NCA knockdown in previously defined sleep-regulatory centers, clock neurons, the visual system or subsets of Dop1R1-expressing neurons did not impact night sleep. FSB: fan-shaped body. MB: mushroom body. Grey and blue box plots: control lines. Red box plots: experimental lines showing reduced night sleep relative to controls. Green box plots: experimental lines failing to show reduced night sleep relative to one or both controls. See Figure 3 figure supplement 1 for n-values and statistical comparisons. (B) Confocal z-stacks of adult male brains expressing genetically-encoded fluorophores under the *A05* or *C01*-Gal4 drivers. Neuropil regions are labelled with anti-Bruchpilot (BRP). Scale bar = 100 µm. Arrows point to neuropil centers. AOTU: anterior optic tubercle. MBNs: mushroom body neurons. OL: optic lobe. AMMC: antennal mechanosensory and motor center. AVP: anterior ventrallateral protocerebrum. SMP: supra-medial protocerebrum. (C-D) *Nca* knockdown in both *A05* and *C01*-neurons recapitulates the effect of pan-neuronal *Nca* knockdown, whereas *Nca* knockdown in either neuronal subpopulation alone did not reduce sleep relative to controls. Mean sleep patterns in 8L: 16D conditions are shown in (C). Median night sleep levels are shown in (D). + > *kk*: n = 80; *C01* > +: n = 64, *C01 > kk*: n = 80; *A05* > +: n = 31; *A05* > *kk*: n = 31; *C01/A05* > +: n = 42; *C01/A05 > kk*: n = 71. *p < 0.05, **p < 0.01, ***p < 0.001, ns – p > 0.05 compared to driver and RNAi alone controls, Kruskal-Wallis test with Dunn’s post-hoc test.

### NCA acts in a common pathway with the Dop1R1 dopamine receptor

In *Drosophila*, dopamine is a pro-arousal factor, with elevated dopaminergic neurotransmission strongly reducing sleep (Kume et al., 2005). Recent studies have shown that the pro-arousal effect of elevated dopamine is mediated by the D1-type Dop1R1 dopamine receptor (Liu et al., 2012; Ueno et al., 2012). Furthermore, both hypo- and hyper-dopaminergic signalling within the human basal ganglia has been proposed to underlie forms of primary dystonia (Breakefield et al., 2008). We therefore tested whether *Nca* promotes sleep in a common pathway with genes involved in dopaminergic signalling. Indeed, we found that heterozygosity for a null or strongly hypomorphic allele of the *Dop1R1* dopamine receptor (*Dop1R1*^MI03085-GFST.2^, a homozygous lethal MiMIC insertion) rescued night sleep loss in *Nca*^*KD*^ flies (Figure 2G, H). Importantly, in both *elav*-Gal4/+ and *kk*/+ control backgrounds, heterozygosity for *Dop1R1*^MI03085-GFST.2^ did not alter sleep levels (Figure 2G, H; p > 0.05, Kruskal-Wallis test with Dunn’s post-hoc test). A similar epistatic interaction between *Nca* and *Dop1R1* was observed using a second, weaker *Dop1R1* allele (*Dop1R1*^MI004437^) (Figure 2 – figure supplement 7).

Mammalian Hippocalcin regulates cell-surface levels of NMDA receptors during LTD (Jo et al., 2010), and *Drosophila nmda receptor 1* (*dNR1*) mutants exhibit reduced sleep during the night (Tomita et al., 2015), similarly to *Nca* knockout and knockdown flies. However, in contrast to *Dop1R1*, we found no signatures of genetic interaction between *Nca* and *dNR1*, suggesting that these loci act in distinct pathways to promote sleep (Figure 2 – figure supplement 8).

### NCA acts in two distinct circuits to promote night sleep

We next sought to delineate the neural circuits in which NCA functions to promote night sleep. Using transgenic RNAi, we performed an extensive screen of sleep relevant circuits defined by numerous *promoter*-Gal4 driver lines (Figure 3A, Figure 3 – figure supplement 1). These include clock neurons, dopaminergic and other neurotransmitter-specific subtypes, fan-shaped body neurons, mushroom body (MB), and sensory neurons (Figure 3A) (Donlea et al., 2011; Joiner et al., 2006; Lamaze et al., 2017; Liu et al., 2014; Pitman et al., 2006; Seidner et al., 2015; Sitaraman et al., 2015). Given the genetic interaction between *Nca* and *Dop1R1*, we also utilised genomic enhancer elements in the *Dop1R1* locus to drive *Nca* knockdown in subsets of potential Dop1R1-expressing neurons (Figure 3A, Figure 3 – figure supplement 1) (Jenett et al., 2012; Jiang et al., 2016). However, in contrast to broadly expressed drivers (*elav*-, *nsyb*- and *inc*-Gal4), *Nca* knockdown in restricted neural subsets was insufficient to significantly reduce night sleep (Figure 3A, Figure 3 – figure supplement 1).

These results suggested a complex sleep-relevant circuit requirement for NCA. We therefore reduced NCA levels in multiple sub-circuits to test for a simultaneous role of NCA in distinct anatomical regions. Through this approach, we found that *Nca* knockdown using two *enhancer*-Gal4 lines (*R14A05* – an enhancer in the *single-minded* locus, and *R72C01* – an enhancer in the *Dop1R1* locus; see Figure 3B for expression patterns) was sufficient to strongly phenocopy the effect of pan-neuronal *Nca* knockdown on night sleep (Figure 3C, D; compare Figure 3C with Figure 2A). For simplicity we refer to these drivers as *A05* and *C01* respectively.

The *A05* enhancer drives expression in approximately 70 neurons, as quantified using a fluorescent nuclear marker (Figure 3 – figure supplement 2A, B) that include a subset of MB neurons, a cluster of cell bodies adjacent to the anterior ventrolateral protocerebrum (AVP), and two visual sub-circuits: optic lobe (OL) and anterior optic tubercle (AOTU) neurons (Figure 3B). *C01* drives expression in approximately 250 neurons (Figure 3 – figure supplement 2C, D) that include the MBs, neurons projecting to the MB γ-lobes, the antennal mechanosensory and motor center (AMMC) (Figure 3B) and the superior medial protocerebrum (SMP). Both drivers label additional cell bodies of unknown identity. The potential overlap of *A05* and *C01* in the MBs raised the possibility that sleep loss in *A05/C01 > Nca* RNAi flies was due to strong NCA knockdown in neurons common to both the *A05* and *C01* enhancers. If so, driving *Nca* RNAi with two copies of either *A05* or *C01* should mimic sleep loss in *A05/C01 > Nca* RNAi flies. However, this was not the case (Figure 4 – figure supplement 3). Thus, NCA is simultaneously required in two non-overlapping sub-circuits defined by the *A05* and *C01* enhancers.

Given that *C01* is a *Dop1R1* enhancer element, that *Nca* and *Dop1R1* genetically interact to regulate sleep (Figure 2G, H), and that Dop1R1 is highly expressed in the MBs (Lebestky et al., 2009), we tested whether the MBs were a constituent of the *C01* expression domain by swapping *C01* for the MB-specific driver *ok107* and measuring sleep in flies expressing *Nca* RNAi in both *A05* and MB neurons. Indeed, knockdown of *Nca* in both *A05* and MB neurons also specifically reduced night sleep (Figure 3 – figure supplement 4), albeit to a weaker degree compared to knockdown in *A05* and *C01* neurons (compare with Figure 3C, D). Thus, we conclude that the MBs are a component of a complex network defined by *C01*- Gal4 with additional, as yet undefined, neurons acting within both the *C01* and *A05* domains to regulate night sleep.

### NCA promotes sleep by suppressing synaptic output from a wake-promoting circuit

We next assessed how NCA impacts excitability of *C01* and *A05* neurons. To do so, we expressed a genetically-encoded fluorescent indicator of neurotransmitter release, UAS-*synaptopHluorin* (*spH*) (Miesenbock, 2012), in *C01* and *A05* neurons of either wild type or *Nca*^KD^ males. spH is localised to presynaptic neurotransmitter-containing vesicles and increases in fluorescence in a pH-dependent manner upon fusion of synaptic vesicles with the presynaptic membrane, providing an optical read-out of neurotransmitter release (Miesenbock, 2012). Intriguingly, we found that *Nca* knockdown significantly enhanced spH fluorescence in the MB α/β-lobes and the AMMC but not in the MB γ-lobe region or the SMP (Figure 4).

**Figure 4.**
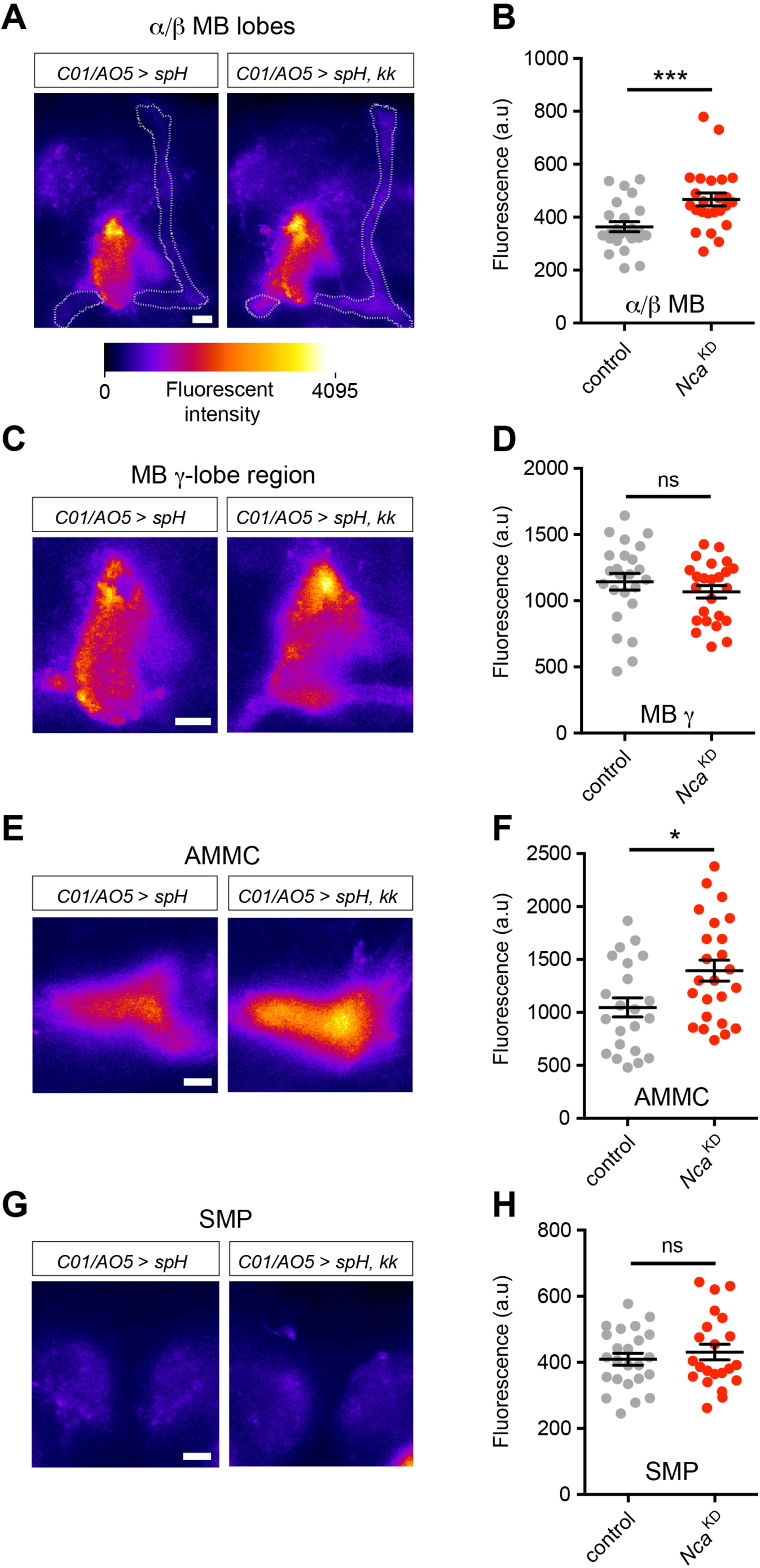
NCA inhibits synaptic release in subsets of *C01*/*A05* neurons. Expression of a fluorescent marker for synaptic release (synapto-pHluorin, spH) in *C01*- and *A05*-positive neurons in control (*C01*/*A05 > spH*) or *Nca* knockdown (*C01/A05 > spH, kk*) backgrounds. Adult brains were dissected and imaged between ZT9-ZT11, when robust sleep loss occurs in *C01*/*A05 > kk* flies (see Figure 3C). (A, C, E, G) Representative pseudo-coloured images for α/β mushroom body lobe Kenyon cells (α/β MB, A; α/β lobes are outlined in white), the MB γ-lobe region (C), AMMC (E) and SMP (G) regions in each genotype. Fluorescent intensity range is shown in the horizontal bar, illustrating minimum to maximum spectrum. Scale bars = 10 μm. (B, D, F, H) Mean fluorescent intensity for individual hemispheres in the above regions. n = 22-24. *p < 0.05, ***p < 0.001, Mann-Whitney U-test.

Since neurotransmitter release is enhanced in subsets of *C01* and *A05* neurons following *Nca* knockdown, this suggested that NCA normally acts to inhibit synaptic output in these circuits. If sleep loss in *Nca* knockdown flies directly results from a loss of such inhibition in *C01* and *A05* neurons (thus enhancing neurotransmitter release), we predicted the following: firstly, that artificial activation of *C01* and *A05* neurons should be sufficient to promote locomotor activity (and thus sleep loss), and secondly, that silencing *C01* and *A05* neurons should suppress sleep loss in *Nca* knockdown flies. To test our first prediction, we enhanced the excitability of *C01* and *A05* neurons by expressing the temperature-sensitive channel TrpA1 in either neuronal subset or both and shifting flies from a non-activating temperature (22°C) to an activating temperature (27°C) sufficient to cause neural excitation through TrpA1-mediated cation influx (Hamada et al., 2008) (Figure 5A). At the non-activating temperature, over-expression of TrpA1 in either circuit or both did not alter sleep levels (Figure 5B). At the activating temperature, excitation of *A05* neurons did not alter night sleep levels relative to controls (Figure 5C, D). In contrast, excitation of *C01* neurons profoundly reduced night sleep (Figure 5C, D) as well as day sleep (Figure 5C). Interestingly, simultaneous activation of *C01* and *A05* neurons further reduced night sleep but not day sleep relative to activation of *C01* neurons alone despite the lack of effect of *A05* neuron activation on sleep (Figure 5C, D).

**Figure 5.**
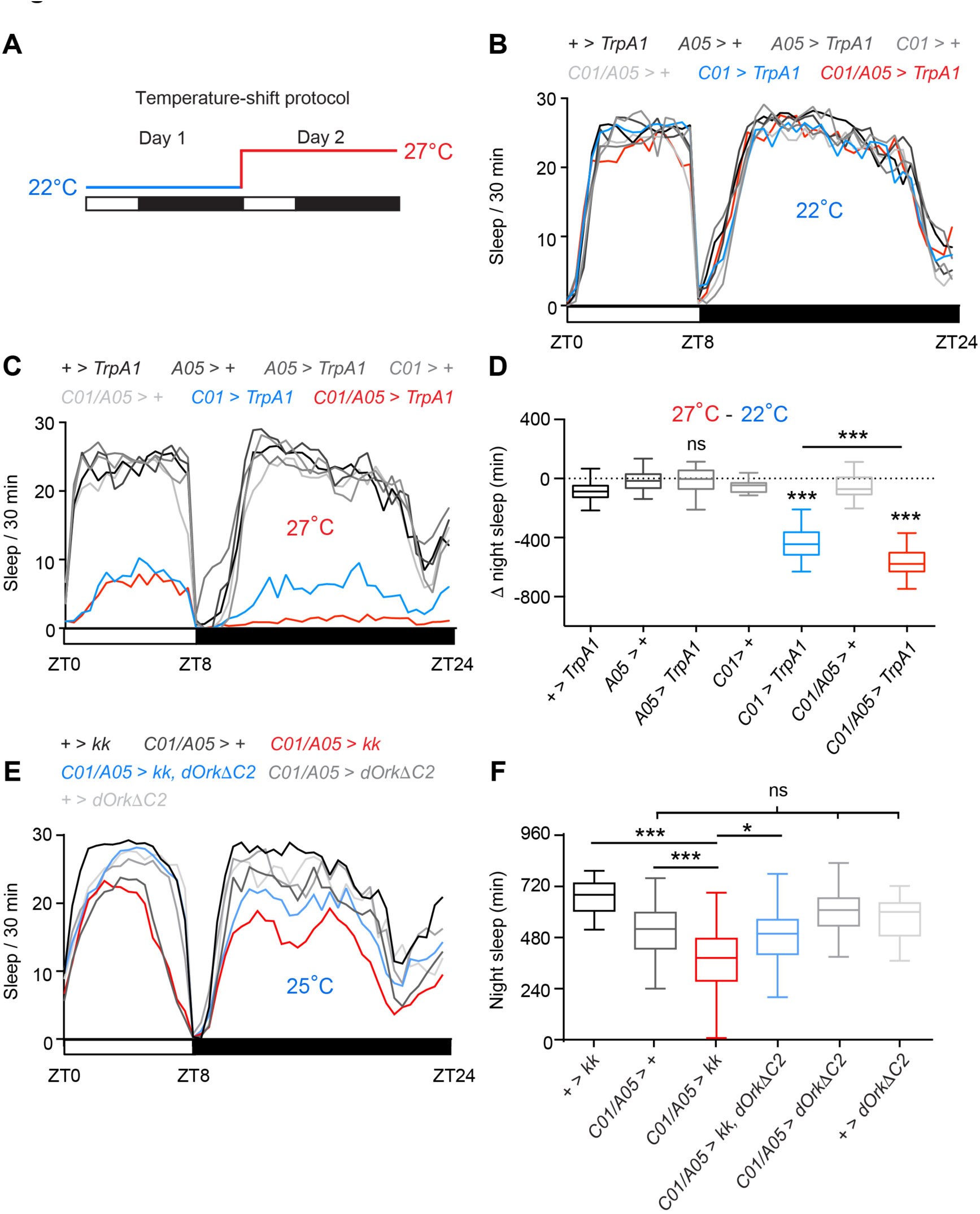
Sleep loss in *Nca* knockdown flies is caused by enhanced excitability of *C01/A05* neurons. (A) Experimental paradigm for acute activation of *A05* or *C01*-neurons. 22°C: non-activating temperature for TrpA1. 27°C: activating temperature. Sleep measurements were measured over two days in 8L: 16D conditions. (B-C) Mean sleep levels across 8L: 16D following expression of TrpA1 in *A05-*, *C01-* or *A05-* and *C01*-neurons (and associated controls) at 22°C (B) or 27°C (C). (D) Median change in night sleep levels (Δ night sleep) following the shift from 22°C on day 1 to 27°C on day 2. + > *TrpA1*: n = 53; *A05* > +: n = 23; *A05 > TrpA1*: n = 68; *C01* > +: n = 24; *C01 > TrpA1*: n = 40; *C01/A05* > +: n = 33; *C01/A05 > TrpA1*: n = 40. ***p < 0.001, ns – p > 0.05, as compared to *TrpA1* or driver alone controls by Kruskal-Wallis test with Dunn’s post-hoc test (for *C01, A05* or *C01/ A05* > *TrpA1* compared to controls) or Mann-Whitney U-test (for *C01/A05 > TrpA1* compared to *C01 > TrpA1*). (E-F) Inhibition of *C01/A05* neurons by expressing dORKΔC2 rescues sleep loss due to *Nca* knockdown in *C01/A05* neurons, while expression of dORKΔC2 does not change base line sleep. Mean sleep patterns in 8L: 16D conditions are shown in (E). Median night sleep levels are shown in (F). *+> kk*: n = 47, *C01/A05 >+*: n = 47, *C01/A05 > kk*: n = 62, *C01/A05 > dORKΔC2, kk*: n = 50, *C01/A05 > dORKΔC2*: n = 39, *+ > dORKΔC2*: n = 30. *p < 0.05, ***p < 0.001, ns – p > 0.05, Kruskal-Wallis test with Dunn’s post-hoc test.

To test our second prediction, we over-expressed a non-inactivating outward rectifying potassium channel (dORKΔC2) in *C01* and *A05* neurons with and without *Nca* knockdown via RNAi. Here, expression of dORKΔC2 is predicted to suppress neuronal firing by hyperpolarizing the resting membrane potential (Nitabach et al., 2002; Park and Griffith, 2006). Silencing *C01* and *A05* neurons with dORKΔC2 in an otherwise wild type background did not alter day or night sleep levels (Figure 5E, F). However, consistent with our above prediction, dORKΔC2 expression significantly suppressed night sleep loss due to *Nca* knockdown in *C01* and *A05* neurons (Figure 5E, F). Thus, we propose that NCA promotes night sleep by limiting synaptic output from a multi-component activity-promoting circuit coordinately defined by subsets of the *C01-* and *A05-*Gal4 expression domains. Our results further suggest that the *C01* and *A05* circuits interact to suppress sleep, with *C01* neurons acting as a predominant pro-arousal circuit and *A05* neurons acting in a modulatory manner to enhance the impact of *C01* activation on night sleep.

## Discussion

Previous work has demonstrated that *Drosophila* Neurocalcin (NCA), a member of the neuronal calcium sensor family, is widely expressed throughout the fly brain and localizes to synaptic regions (Teng et al., 1994). However, the neurobiological functions of NCA have remained unclear. Here we define a role for NCA in promoting night sleep and show that NCA acts via inhibiting synaptic output from a complex movement-promoting circuit.

Previous genetic screens have identified an array of sleep-promoting factors in *Drosophila* (Tomita et al., 2017). However, despite extensive circuit analyses, the activity of such proteins does not fully map onto known sleep-regulatory neurons, and the complete neural substrates in which these factors act have yet to be determined (Afonso et al., 2015; Rogulja and Young, 2012; Shi et al., 2014; Stavropoulos and Young, 2011; Tomita et al., 2015; Wu et al., 2014). Our results are consistent with these findings and offer a tentative explanation for the difficulties in defining circuit requirements for sleep-relevant proteins. We show that NCA activity is not required within a single cell-type or neuropil region to promote sleep. Instead, sleep-relevant NCA activity can largely be localised to *two* distinct domains of the *Drosophila* nervous system defined by the *A05*- and *C01*-Gal4 drivers. One relevant cell-type within these domains is the mushroom bodies (MBs), a known sleep-regulatory center (Joiner et al., 2006; Pitman et al., 2006; Sitaraman et al., 2015). However, the MBs only partially contribute to one of the two sleep-relevant domains, indicating that NCA acts in a dispersed network that likely incorporates several non-overlapping wake-promoting circuits.

Interestingly, whereas activation of *C01* neurons is sufficient to suppress sleep, activation of *A05* neurons reduces night but not day sleep only in the context of

*C01* neuron activation. We note that activation of *C01* neurons profoundly reduces sleep during both day and night periods, whereas only night sleep is impaired in flies with reduced NCA expression in *C01* and *A05* neurons. Based on these findings, we hypothesize that reduction of NCA in *C01* and *A05* neurons causes mild hyperexcitation, which in *C01* neurons alone is insufficient to modulate sleep but in both *C01* and *A05* neurons simultaneously causes an increase in network excitability sufficient to reduce night sleep. A selective inhibition of neurotransmitter release by NCA in subsets of *C01* and *A05* neurons is supported by our in vivo imaging data showing that *Nca* knockdown increases synaptic output in the MB α/β-lobes and AMMC but not the MB γ-lobe region or the SMP. Since NCA is only partly required in the MBs, we consider it unlikely that increased synaptic output within the MBs and AMMC alone is sufficient to drive sleep loss in *Nca* knockdown flies. Rather, we propose that coincident hyperactivity of several sub-circuits within the *C01* and *A05* domains collectively drives night sleep loss. Temporal information encoded by light-sensing or circadian pathways may further demarcate the period in which NCA promotes sleep (i.e the night), with such inputs likely to be bypassed by ectopic activation of *C01* neurons, resulting in sleep loss across 24 h.

Since silencing of *C01* and *A05* neurons does not alter basal sleep, we suggest that these circuits are normally activated by an ethological stimulus absent during our experimental conditions. We further postulate that modulatory dopamine signalling through Dop1R1 promotes neural activity in wake-promoting *C01* neurons. These neurons are defined by a *Dop1R1* enhancer element, and thus it is likely that at least a subset of *C01* neurons express Dop1R1. Indeed, dopamine signalling via Dop1R1 has been shown to enhance synaptic transmission between antennal lobe projection neurons and MB neurons (Ueno et al., 2013). This model is attractive since reducing such input could balance the increased synaptic release caused by *Nca* knockdown, providing an explanatory basis for the genetic interaction between *Nca* and *Dop1R1*.

How might NCA inhibit synaptic output from *C01* and *A05* neurons? The mammalian NCA homolog Hippocalcin acts pleiotropically in several pathways that control neuronal excitability and plasticity, facilitating NMDA receptor endocytosis during LTD and gating the slow afterhyperpolarisation, a calcium-activated potassium current controlling spike frequency adaptation that is thought to be mediated by a complex array of potassium channels (Andrade et al., 2012; Jo et al., 2010; Tzingounis et al., 2007). Recent data suggest that Hippocalcin also negatively regulates calcium influx through N-type voltage-gated calcium channels (Helassa et al., 2017). Given the strong homology between Hippocalcin and NCA, it is possible that NCA suppresses neurotransmitter release through similar pathways in *Drosophila* neurons, although the lack of genetic interaction between *Nca* and the *dNR1* NMDA receptor suggests that an enhancement of NMDA receptor levels is unlikely to significantly contribute to sleep loss in *Nca* knockout/knockdown flies.

Mutations in Hippocalcin also cause the movement disorder DYT2 primary isolated dystonia (Charlesworth et al., 2015). What is the common neurobiological link between dystonic movements in humans and sleep loss in *Drosophila*? We suggest that both phenotypes fundamentally reflect a dysregulation of motor control i.e when activity is initiated, and for how long such activity is maintained. Hippocalcin/Neurocalcin homologs have the potential to act as molecular toggle switches, inhibiting sustained rapid-firing and neurotransmitter release in activity-promoting neurons via simultaneous modulation of potassium and calcium channel function (Helassa et al., 2017; Tzingounis et al., 2007). Mutations in these calcium sensors may thus result in prolonged bouts of synaptic output and corresponding motor activity. We note that *insomniac*, another sleep-promoting gene in *Drosophila*, is homologous to the myoclonus dystonia-gene *KCTD17* (as well the paralogs *KCTD2* and *KCTD5*) (Li et al., 2017; Mencacci et al., 2015; Stavropoulos and Young, 2011). It will thus be intriguing to test if other dystonia gene homologs also promote sleep in *Drosophila,* and whether human dystonia and *Drosophila* sleep loss can thus be considered homologous phenotypes (or phenologs) linked to mutations in a conserved genetic network that functions to suppress inappropriate activity in both humans and flies (Lehner, 2013; McGary et al., 2010).

## Materials and Methods

### Fly husbandry

Flies were maintained on standard fly food at constant temperature 25°C under 12 h: 12 h light-dark cycles (12L: 12D). The following strains were obtained from the Bloomington and VDRC stock centers: kk108825 (v100625), hmj21533 (54814), jf03398 (29461), *Dop1R1*^MI03085-GFSTF.2^ (59802), *Dop1R1*^MI04437^(43773), *ple*-Gal4 (8848), *Chat*-Gal4 (6798), *vGlut*-Gal4 (26160), *GAD*-Gal4 (51630), *Ddc*-Gal4(7010), *GMR*-Gal4 (1104), *Trh.1*-Gal4 (38388), *Tdc2*-Gal4(9313), *C5*-Gal4 (30839), *ok107*-Gal4 (854) and *A502* (16130). The remaining lines obtained from the Bloomington stock center are part of the Janelia Flylight collection with identifiable prefixes: R23E10-Gal4, R55B01-Gal4, R52H12-Gal4, Hdc-Gal4 (R17F12-Gal4), R14A05-Gal4, R72B05-Gal4, R72B07-Gal4, R72B08-Gal4, R72B11-Gal4, R72C01-Gal4, and R72C02-Gal4. The following lines were gifts from laboratories of Kyunghee Koh: *elav*-Gal4, *nsyb*-Gal4, *tim*-Gal4 and *TUG*-Gal4; Joerg Albert: *nompC*-Gal4 (Kamikouchi et al., 2009) and Nicolas Stavropouplos: *inc*-Gal4:2 (Stavropoulos and Young, 2011). *ppk*-Gal4 and *TrpA1*-CD-Gal4 were described previously (Zhong et al., 2012). *GMR-hid*, *tim*^*KO*^ and *cry*^*02*^ were previously described in (Lamaze et al., 2017). Except for *Ddc*-Gal4, *Trh.1*-Gal4, *Tdc2*-Gal4, *nompC*-Gal4 and *Hdc*-Gal4, all *Drosophila* strains above were either outcrossed five times into an isogenic control background (*iso31*) or insertion-free chromosomes were exchanged with the *iso31* line (*hmj21533, jf03398, Dop1R1^MI03085-G^*, *Dop1R1*^*MI04437*^ and *NMDAR1*^MI11796^) before testing for sleep-wake activity behaviour. Note: R14A05-Gal4 was initially mislabelled as R21G01-Gal4 in Bloomington shipment. The clear mismatch between the image of R21G01>GFP in FlyLight database and our immunostaining data (A05, Fig3B) led us to clarify the identity of the line, as R14A05-Gal4, by sequencing genomic PCR product using primers pair: pBPGw_ampF: agggttattgtctcatgagcgg and pBPGw_Gal4R: ggcgcacttcggtttttctt.

### Generation of the *Nca*^*KO*^ line

Null alleles of *Nca* were generated using homologous recombination as described previously (Baena-Lopez et al., 2013). Briefly, genomic DNA was extracted from 20 wild type flies (Canton S) using the BDGP buffer A-LiCl/KAc precipitation protocol (http://www.fruitfly.org/about/methods/inverse.pcr.html). The 5’ (Arm 1) and 3’ (Arm 2) genomic regions flanking the *Nca* coding sequence were PCR amplified via high fidelity DNA polymerase (Q5 high-fidelity 2X master mix, M0492S, NEB) with the following primers: NotI_Arm1F1:gcggccgctaatttgcagctctgcatcg, NotI_Arm1R1:gcggccgcatggtaagaagcacgcaacc, AscI_Arm2F1:ggcgcgccttatgaccgttccaaaacacc, AvrII_Arm2R1:cctaggggctaaatacgttgaccaagc. The corresponding Arm1 and Arm2 fragments (~2.5kb) were gel purified (Wizard^®^ SV Gel and PCR Clean-Up System, A9281, Promega) and cloned into pCR-Blunt II-TOPO vector (Zero Blunt^®^ TOPO^®^ PCR Cloning Kit, 450245, ThermoFisher Scientific), and subsequently sub-cloned via NotI (R3189S, NEB) and AscI/AvrII digestion (R0558S and R0174S, NEB) and T4 ligation (M0202S, NEB) into pTV vector, a P-element construct containing the mini-*white*^+^ marker and UAS-*reaper* flanked by FRT and I-SceI sites (Baena-Lopez et al., 2013). The sequence identifies of Arm 1 and Arm 2 fragments within the pTV vector were verified via Sanger sequencing using the following primers: nca1_f: cagctctgcatcgctttttgt, nca1_3_f: ccctcgcgcatggtacttta, nca1_r: agcgtcacataagttctccca, nca1_4_f: tggacgaaaataacgatggtca, nca1_5_f: agactacttagccatgttttcatact, nca1_2_f: tgacgaagccacaattaaagagtg, nca1_1_f: gcaaccctgttcccctttca, nca2_f: gaccgttccaaaacaccca, nca2_3_f: ttgttgtgcgccacgttttc, nca2_r: acgtatgctccatgattcctct nca2_4_f: tgcaggtcggttaatcaatgc, nca2_5_f: tcaatcgatttggggccagg, nca2_2_f: ccttctccaggctcagcaaa, nca2_1_f: actctgcatttcgataagattagcc. Donor lines containing pTV vector with Arm1 and Arm2 homologous fragments (pTV_nca1+2) were then generated via embryonic injection and random P-element mediated genomic insertions (Bestgene, inc.). To initiate homologous recombination between pTV_nca1+2 and the endogenous *Nca* locus, donor lines were crossed to *yw*; hs-*flp*, hs-I-*SceI*/CyO and the resulting larvae were heat shocked at 48 h and 72 h after egg laying for 1 h at 37°C. Around 200 female offspring with mottled/mosaic red eyes were crossed in pools of three to *ubiquitin*-Gal4[3xP3-GFP] males to remove nonspecific recombination events (via UAS-*reaper*-mediated apoptotic activity). The crossings were flipped once over and the progeny (~12000 adults) was screened for the presence of red-eyed and GFP-positive flies. Three independent GFP^+^ red-eyed lines (*ko1*, *ko2*, and *ko3*) were identified. The exchange of endogenous *Nca* locus with pTV_nca1+2 fragments was confirmed by detecting a 2.6 kb PCR product (Figure 1 figure supplement 1C) in the genomic DNA samples of the above three lines (pre-digested by EcoRI/NotI) using the following primer pairs: ncaKO-F2: tgggaattgactgatacagcct; ncaKO-R2: ggcactacggtacctgcat. ncaKO-F2 matches to the region between 24 bp and 2 bp upstream of Arm1 and ncaKO-R2 overlaps with attP site (Figure 1A). The absence of endogenous *Nca* mRNA in *ko1* flies was confirmed by standard and quantitative RT-PCR (Figure 1 figure supplement 1D; see also the below RNA section). The min-*white*^+^ cassette and majority of pTV vector sequences were further removed from the *ko1* genome via Cre-loxP recombination (Figure 1A). This “Cre-out” strain was then backcrossed five times to a *Nca*^*A502*^ line (where *A502* is a P-element insertion 2 kbp upstream of the *Nca* CDS) that was outcrossed previously into the *iso31* background (see Fly husbandry section). Before testing for changes in sleep/wake behaviour, the resulted line, termed *Nca* knockout (*Nca*^KO^), was lastly verified by sequencing a 576 bp genomic PCR product (using primer pair: nca1_5_f and nca2_r), confirming the absence of *Nca* CDS sequence and the insertion an attP site in the *Nca* locus.

### RNA extraction and Quantitative PCR

For RNA extractions, 10-20 fly heads per genotype were collected with liquid nitrogen and dry ice. Total RNA was extracted using TRIzol™ reagent following manufacturer’s manual (Thermo Fisher Scientific). cDNA was reverse transcribed from 250 or 500 ng of DNase I (M0303S, NEB) treated RNA via MMLV RT (M170A, Promega). A set of five or six standards across 3125-fold dilution was prepared from the equally pooled cDNA of all genotypes in each experiment. Triplicated PCR reactions were prepared in 96-well or 384-well plates for standards and the cDNA sample of each genotype (20 to 40 fold dilution) by mixing in Power SYBR Green Master Mix (Thermo Fisher Scientific) and the following primer sets: ncaqF2: acagagttcacagacgctgag, ncaqR2: ttgctagcgtcaccatatggg; cg7646F: gcctttcgaatgtacgatgtcg, cg7646R: cctagcatgtcataaattgcctgaac or rp49F:cgatatgctaagctgtcgcaca, rp49R: cgcttgttcgatccgtaacc. PCR reactions were performed in Applied Biosystems StepOne (96-wells module) or QuantStudio 6Flex instruments (384 wells module) using standard thermocycle protocols. Melting curve analysis was also performed to evaluate the quality of the PCR product and avoid contamination. The Ct values were exported as csv files and a standard curve between Ct values and logarithm of dilution was calculated using the liner regression function in Graphpad. The relative expression level for *Nca*, *cg7646* and *rp49* of each sample were estimated by interpolation and anti-logarithm. The expression levels of *Nca* and *cg7646* for each genotype were further normalized to their respective average *rp49* expression level. Statistical differences between the normalized expressions levels of each genotype were determined by Mann-Whitney test or Kruskal-Wallis test with Dunn’s post-hoc test using Graphpad software.

### Sleep-wake behavioral analysis

Three to five days old male or virgin female flies were collected and loaded into glass tubes containing 4% sucrose and 2% agar (w/v). Sleep-wake behavior was recorded using the *Drosophila* Activity Monitor (DAM, TriKinetics, inc.) system or *Drosophila* ARousal Tracking (DART, BFKlab) system for 3 days in the designated LD regime (L12: D12 or L8: D16) at 25°C. Behavioral recordings from the third day of the given LD regime were then analyzed. All flies were entrained to 12L: 12D prior to entering designated LD regimes. For ectopic activation experiments involving UAS-*TrpA1*, flies were cultured in 18°C during development and then entrained to L8: D16 at 22°C before entering L8: D16 condition at 27°C. *Drosophila* activity (or wake) is measured by infra-red beam crosses in DAM or by direct movement tracking in DART. Sleep is defined by 5 minutes of inactivity (where inactivity is defined as no beam crosses during 1 min in the DAM or less than 3 mm movement in 5 s in DART). The csv output files with beam crosses (DAM) or velocity data (DART) were processed by a customized Excel calculators (Supplementary file 1) and R-scripts (https://github.com/PatrickKratsch/DAM_analysR) to calculate the following parameters for individual flies: *Onset and offset of each sleep bout*, *sleep bout length*, *day and night sleep minutes*, *daily total sleep minutes,* and *daily sleep profile* (30 minutes interval). An established MATLAB^®^ based tool, Flytoolbox, was used for circadian rhythmicity analysis (Levine et al., 2002a, b). Briefly, the strength of rhythmicity (RI) was estimated using the height of the third peak coefficient in the auto-correlogram calculated for the activity time series of each fly. Rhythmic Statistics values were then obtained from the ratio of the RI value to the 95% confidence interval for the correlogram (2/√N, where N is the number of observations, which correlatively increase with the sampling frequency), in order to determine statistical significance of any identified period (RS is ≥ 1)

### Immunohistochemistry and confocal microscopy

Adult male R72C01 > CD8::*GFP* and R21G01 > CD4::*tdTomato* flies were anesthetized in 70% ethanol before brains were dissected in PBT (0.1M phosphate buffer with 0.3% TritonX100) and collected in 4% paraformaldehyde/PBT on ice. The fixation was then performed at room temperature for 15 min before washing 3 times with PBT. The brain samples were blocked using 5% goat serum/PBT for 1 h at room temperature before incubation with primary antibodies. The samples were washed 6 times with PBT before incubated with Alexa Fluor secondary antibodies in 5% goat serum/PBT at 4°C over 24 h. After washing 6 times with PBT, the samples were mounted in SlowFade Gold antifade reagent (S36936, Thermo Fisher Scientific) on microscope slides and stored at 4°C until imaged using an inverted confocal microscope Zeiss LSM 710. Primary antibody concentrations were as follows: mouse anti-nc82 (Developmental Studies Hybridoma Bank) - 1:200; rabbit anti-GFP (Invitrogen) - 1:1000; rabbit anti-dsRED (Clontech) - 1:2000. Alexa Fluor secondaries (Invitrogen) were used as follows: Alexa Fluor 647 goat anti-mouse IgG - 1:500, Alexa Fluor 488 goat anti-rabbit IgG - 1:2000, Alexa Fluor 555 goat anti-rabbit IgG - 1:2000. For quantification of nuclei number in *C01 > red-stinger* and *A05 > red-stinger* brains, unstained Red-Stinger fluorescence was captured via confocal microscopy. DAPI (Sigma Aldrich) was used to counterstain nuclei (at a dilution of 1:5000). The number of Red-Stinger-positive nuclei in each brain was subsequently quantified using the ImageJ 3D Objects Counter tool, with a variable threshold used to incorporate all of the visible Red-Stinger-positive nuclei.

### Synapto-pHluorin imaging

Synaptic activity of C01/A05 neurons was monitored in *ex vivo* fly brains using UAS- super-eclipse-synaptopHluorin construct (UAS-*spH*) (Miesenbock, 2012). Adult male *C01/A05 >* UAS-*spH* or *C01/A05 >* UAS-*spH, kk* flies were housed in normal behaviour tubes (see behaviour analysis section) and entrained for 3 days in L8: D16 condition at 25°C. Individual flies of either genotype were carefully captured between ZT9 and ZT11 and fly brains were immediately dissected in HL3 *Drosophila* saline (70 mM NaCl, 5 mM KCl, 1.5 mM CaCl_2_, 20 mM MgCl_2_, 10 mM NaHCO_3_, 5 mM Trehalose, 115 mM Sucrose and 5 mM HEPES, pH 7.2) at room temperature. Fly brains were transferred into 200 μl HL3 in a poly-lysine treated glass bottom dish (35 mm, 627860, Greiner Bio-One) before imaging using an inverted confocal Zeiss LSM 710 microscope (20x objective with maximum pinhole). Three to five image stacks (16 bits) were taken within two minutes to minimise tissue degradation and to cover the depth of all spH-positive anatomical regions. Z-projections of the image stacks of each brain were generated by ImageJ software before the fluorescent intensity of the indicated neuropil centres was quantified using free drawn ROIs. Background fluorescence measured by the same ROIs from areas with no brain tissue was then subtracted to obtain the final fluorescent value. Mean fluorescent values of the indicated neuropil regions in each hemisphere were calculated and compared between genotypes. The statistical difference was determined by Mann-Whitney U-test using Graphpad software.

### Bioinformatics

Conservation of amino acid residues between *Drosophila* Neurocalcin and human Hippocalcin was determined using ClustalW2 software for multiple sequence alignment. Amino-acid identity and similarity was visualised using BOXSHADE.

## Acknowledgements

We thank Jack Humphrey for performing initial work on *Neurocalcin* knockdown flies, and Kyunghee Koh and Simon Lowe for helpful comments on the manuscript. This study was supported by the Wellcome Trust (Synaptopathies strategic award [104033]), and by the MRC [New Investigator Grant MR/P012256/1]. P.K is supported by a Wellcome Trust Neuroscience PhD studentship.

## Footnotes

**Competing interests** The authors have no financial or non-financial competing interests.

## Figures and Figure legends

**Figure 1 supplemental figure 1.**
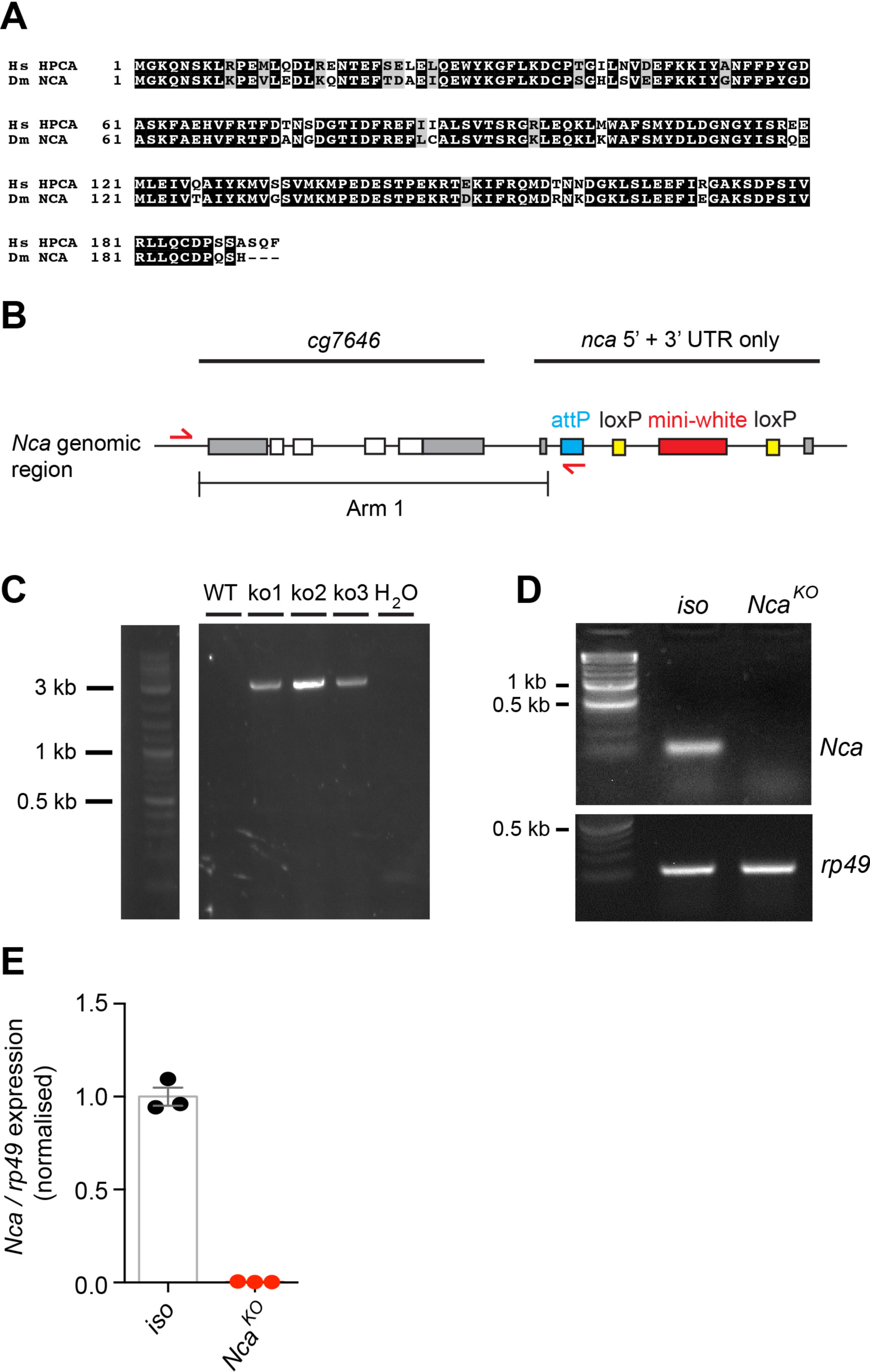
Generation of the *Nca* knockout allele. (A) Hippocalcin and Neurocalcin are highly conserved neuronal calcium sensors. Amino-acid alignment of human (Hs) Hippocalcin (HPCA) and *Drosophila* (Dm) Neurocalcin (NCA). Identical amino-acids are shaded in black, with functionally similar amino-acids shaded in grey. Total homology between Hippocalcin and Neurocalcin is > 90%. (B-C) PCR validation of homologous recombination events. Correct recombination was verified using primers designed to the attP site and upstream of the neighbouring locus *cg7646* (B), which will only generate a ~ 3 kb product following homologous recombination between the targeting vector and the *Nca* locus (C). Three independent targeting events (ko1-3) are shown in (C), one of which was selected for mini-*white* removal as described in Figure 1. WT: wild-type genome lacking an attP site neighbouring the *cg7646* locus. (D-E) No *Nca* mRNA was detected in *Nca*^KO^ using either standard RT-PCR (D) or quantitative RT-PCR (E; n = 3 qPCR reactions for *iso31* control and *Nca*^KO^ flies).

**Figure 1 supplemental figure 2.**
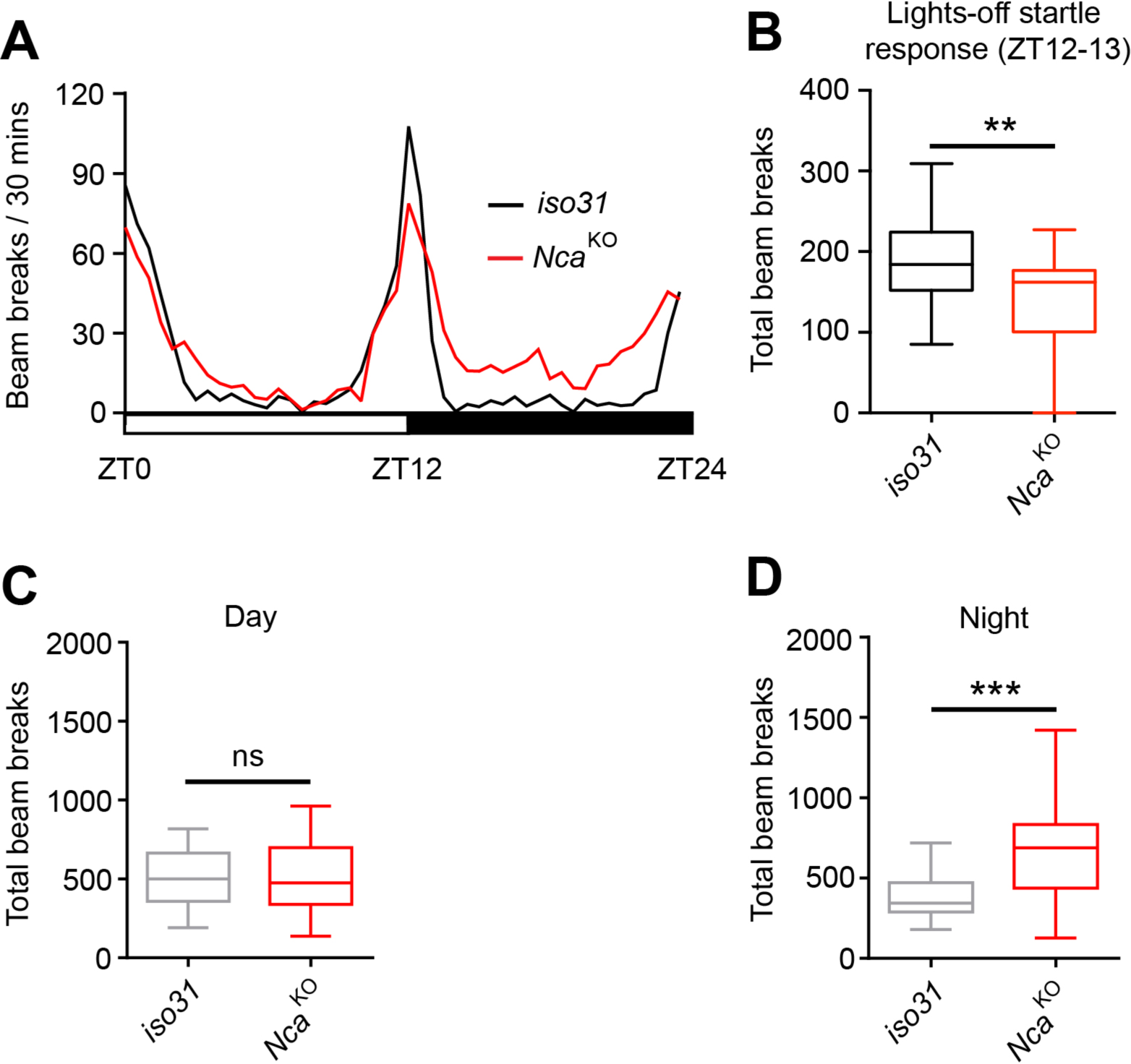
Reduced startle response and elevated night time locomotor activity in *Nca*^KO^ flies. (A) Mean level of locomotor activity (beam crosses) for *iso31* control and *Nca*^KO^ males under 12L: 12D conditions. (B) Median beam breaks during the hour immediately following lights-off (ZT12-13). The rapid light-dark transition initiates a startle response and thus elevated locomotor activity, the degree of which is reduced in *Nca*^KO^ males. (C-D) Median total beam crosses during the day (C) and night (D) periods. Locomotor activity is significantly increased in *Nca*^KO^ males during the night but not the day. n = 32 per genotype. **p < 0.01, ***p < 0.001, ns – p > 0.05, Mann-Whitney U-test.

**Figure 1 supplemental figure 3.**
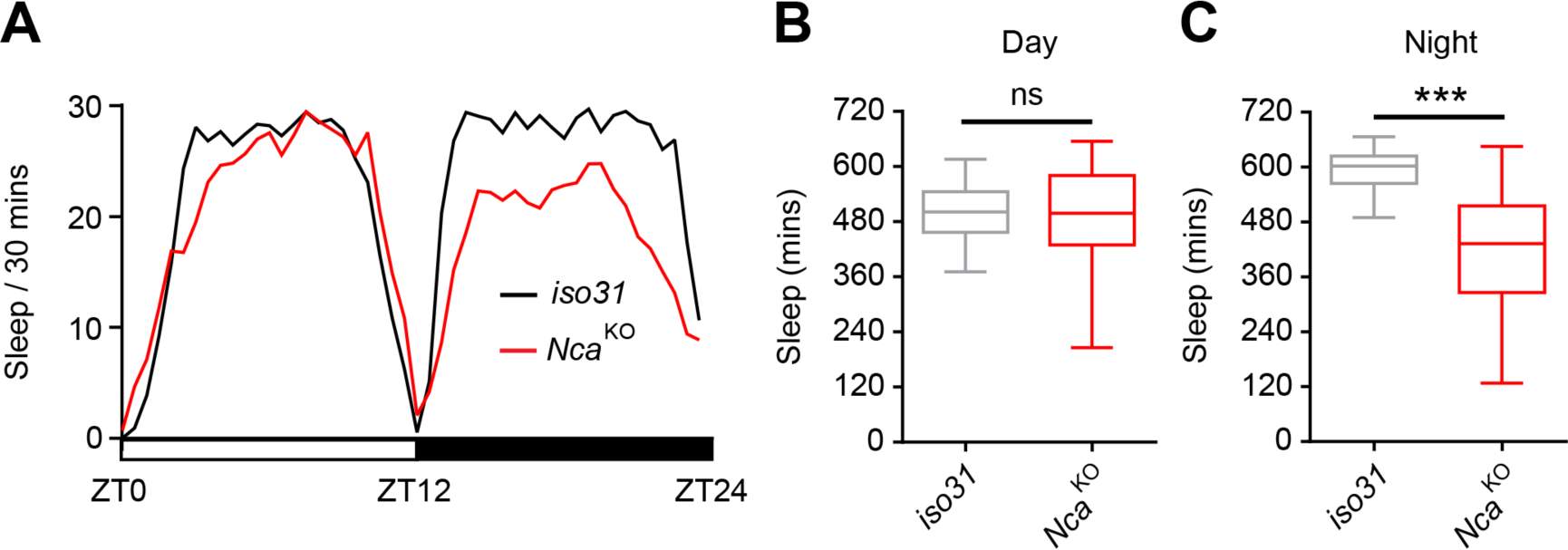
Night time sleep loss in *Nca*^KO^ flies. (A) Mean sleep level of *iso31* control and *Nca*^KO^ flies under 12L: 12D conditions. (B-C) Median total day (B) and night (C) sleep in *iso31* control and *Nca*^KO^ flies. Night sleep is significantly reduced in *Nca*^KO^ flies compared to controls. n = 32 per genotype. ***p < 0.001, ns – p > 0.05, Mann-Whitney U-test.

**Figure 1 supplemental figure 4.**
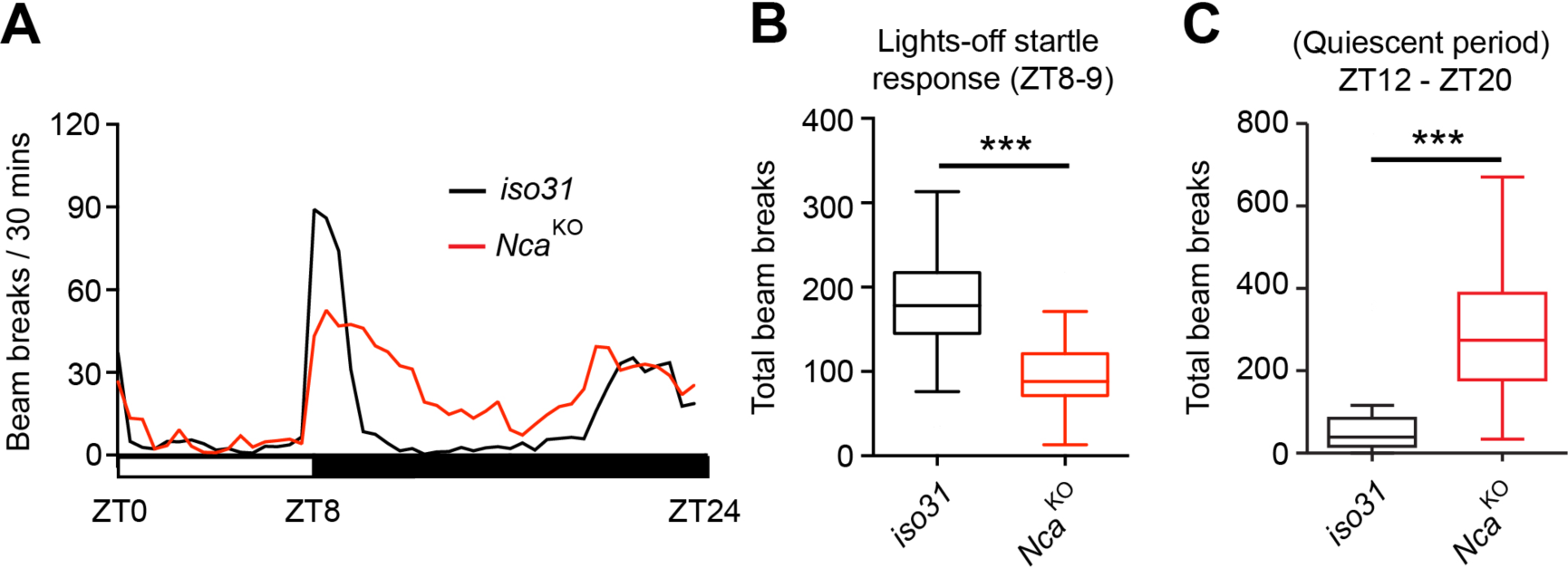
Reduced startle response and elevated night time locomotor activity in *Nca*^KO^ flies under 8L: 16D conditions. (A) Mean level of locomotor activity (beam crosses) for *iso31* control and *Nca*^KO^ males. (B) Median beam breaks during the hour immediately following lights-off (ZT8-9). (C-D) Median total beam crosses during the normally quiescent period of the night (ZT12-20). n = 16 per genotype. ***p < 0.001, Mann-Whitney U-test.

**Figure 1 supplemental figure 5.**
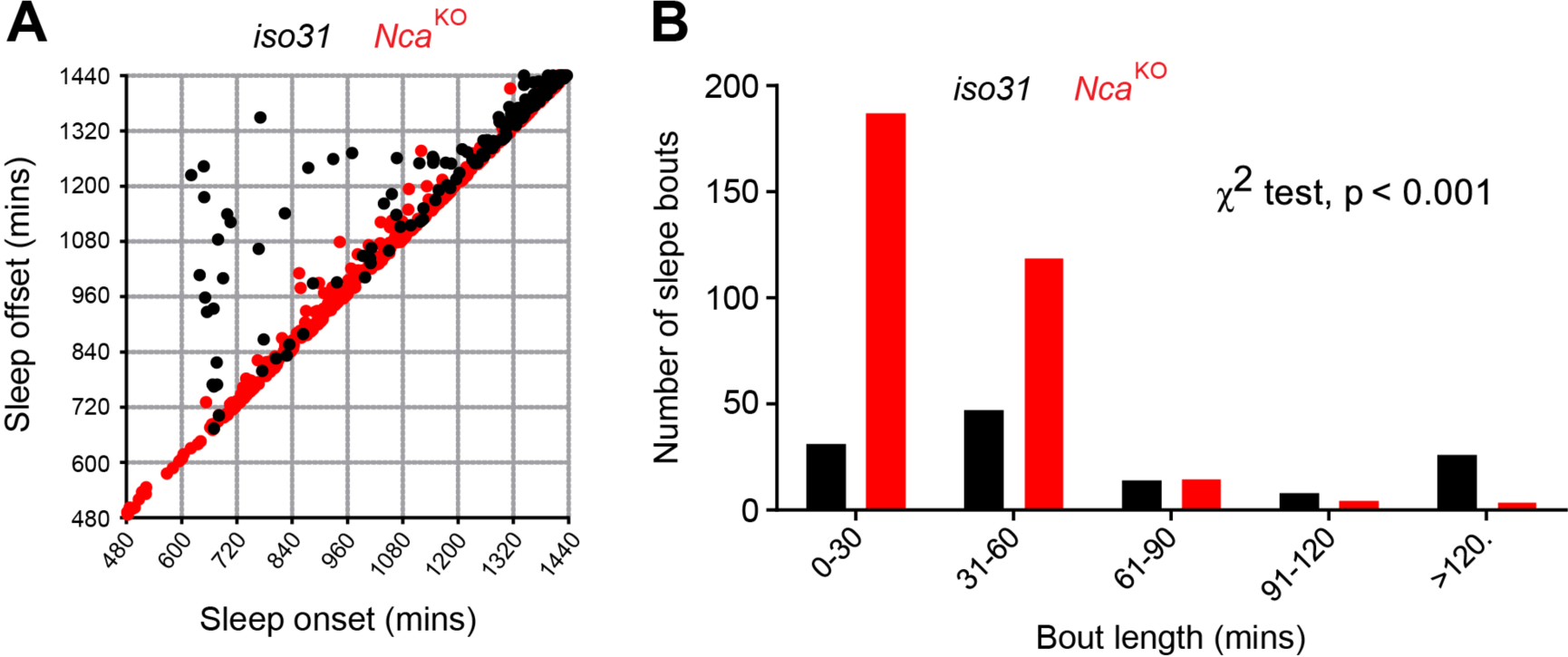
Reduced long sleep bouts in *Nca*^KO^ flies. (A) Following detailed fly movement detection using the DART system, individual sleep bout durations were further estimated using a custom-made R program and visualised by plotting sleep bout offset against onset for night sleep bouts in control *iso31* and *Nca*^KO^ adult males under 8L: 16D conditions. In control flies, longer sleep bouts initiated early during the night (note the black dots deviating from the diagonal), which are largely absent in *Nca*^KO^ adult males (red dots). n = 16 for each genotype. (B) Distribution of sleep bout lengths in *Nca*^KO^ and control adult males. Note the significant shift towards shorter sleep bout lengths in *Nca*^KO^ flies (*Nca*^KO^ vs. *iso31* control: χ^2^, *df*: 85.59, 4, p < 0.001).

**Figure 1 supplemental figure 6.**
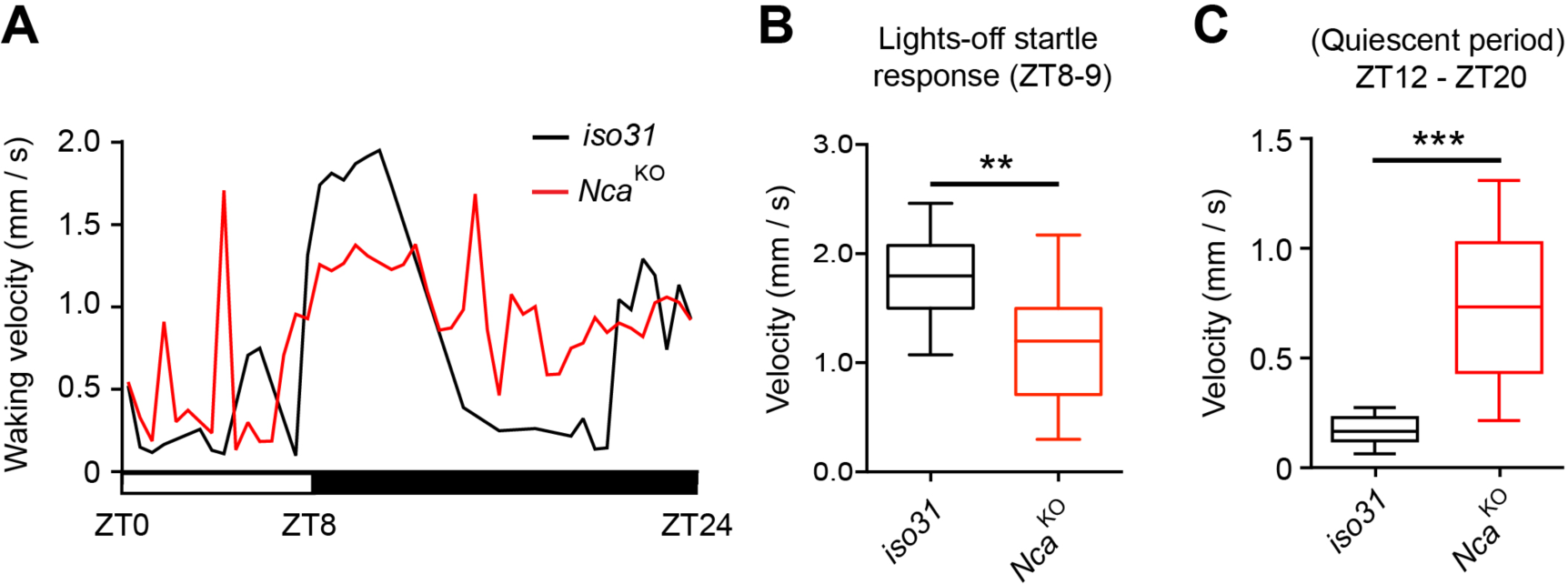
Detailed tracking reveals increased locomotor velocity during the night and reduced velocity during the startle response to lights-off in *Nca*^KO^ flies. (A) Mean waking locomotor velocity in *iso31* control and *Nca*^KO^ adult males under L8: 16D condition. (B) Locomotor velocity is significantly reduced in *Nca*^KO^ flies following the startle response to lights-off (ZT8-ZT9). (C) *Nca*^KO^ flies exhibit a significant increase in locomotor velocity between ZT12-ZT20 compared to controls – a normally quiescent period. n = 16 per genotype. **p < 0.01, ***p < 0.001, Mann-Whitney U-test.

**Figure 2 supplemental figure 1.**
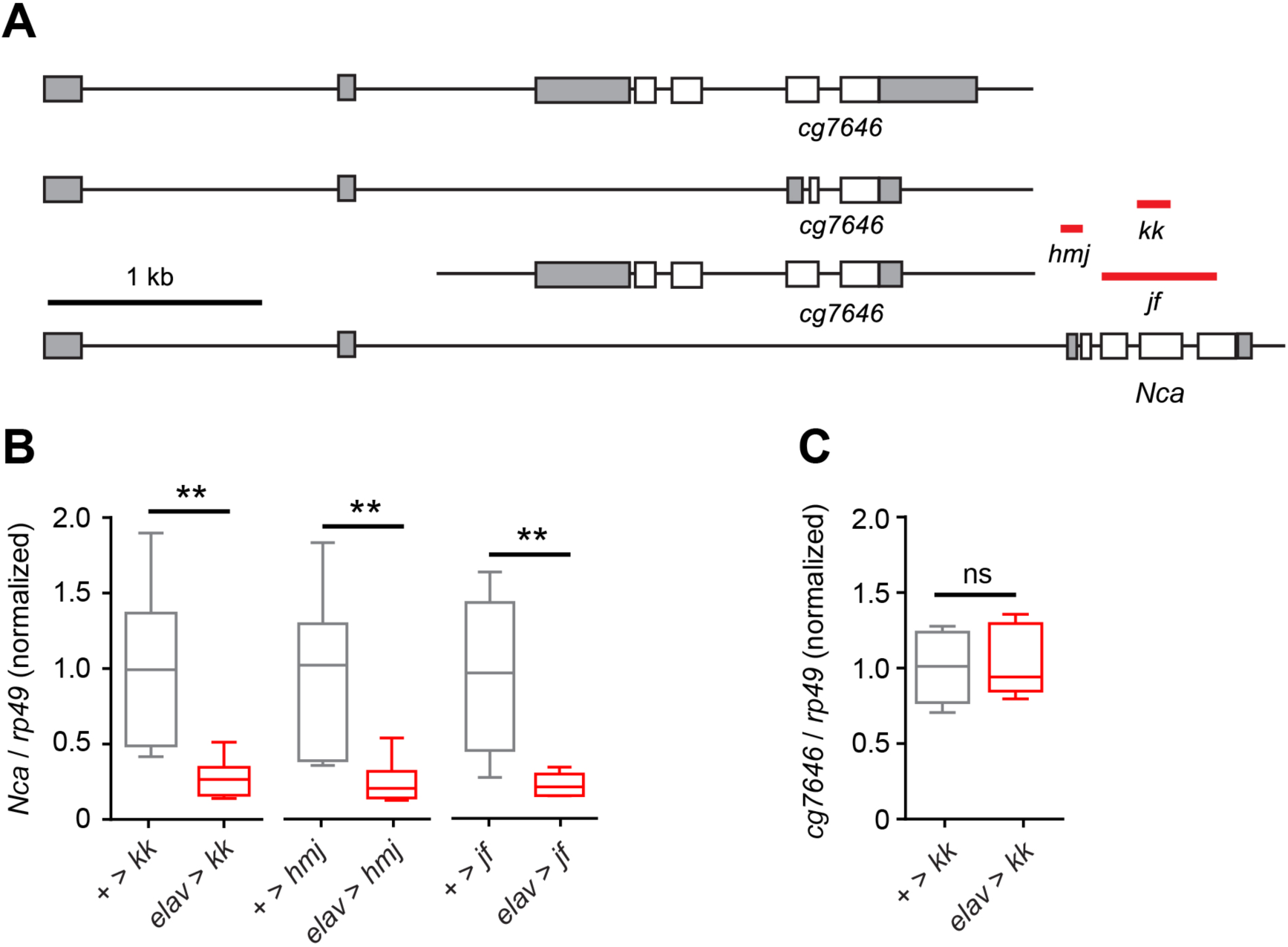
Robust knockdown of *Nca* using independent *elav*-Gal4-driven *Nca* RNAi lines. (A) Schematic showing the *Nca* locus alongside the neighbouring *cg7646* locus, which shares a common promoter region with *Nca*. Regions of the *Nca* transcript targeted by the *kk108825*, *hmj21533* and *jf03398* RNAi lines (termed *kk*, *hmj* and *jf* respectively) are depicted by red bars. (B) qPCR verification of *Nca* knockdown by *kk*, *hmj* and *jf*. RNAi. Transgene insertions lacking the *elav*-Gal4 driver were used as controls. (C) Knockdown of *Nca* had no effect on transcription of the neighbouring *cg7646* locus. Expression levels of *Nca* or *cg7646* were normalised to the *rp49* control transcript and are displayed as the ratio to the mean level of the respective RNAi alone controls (+ > *kk*, + > *hmj* or + > *jf*). n = 6 for all qPCRs (two independent biological repetitions of RNA extraction with triplicated qPCR reactions for each genotype). **p < 0.01, ns – p > 0.05, Mann-Whitney U-test.

**Figure 2 supplemental figure 2.**
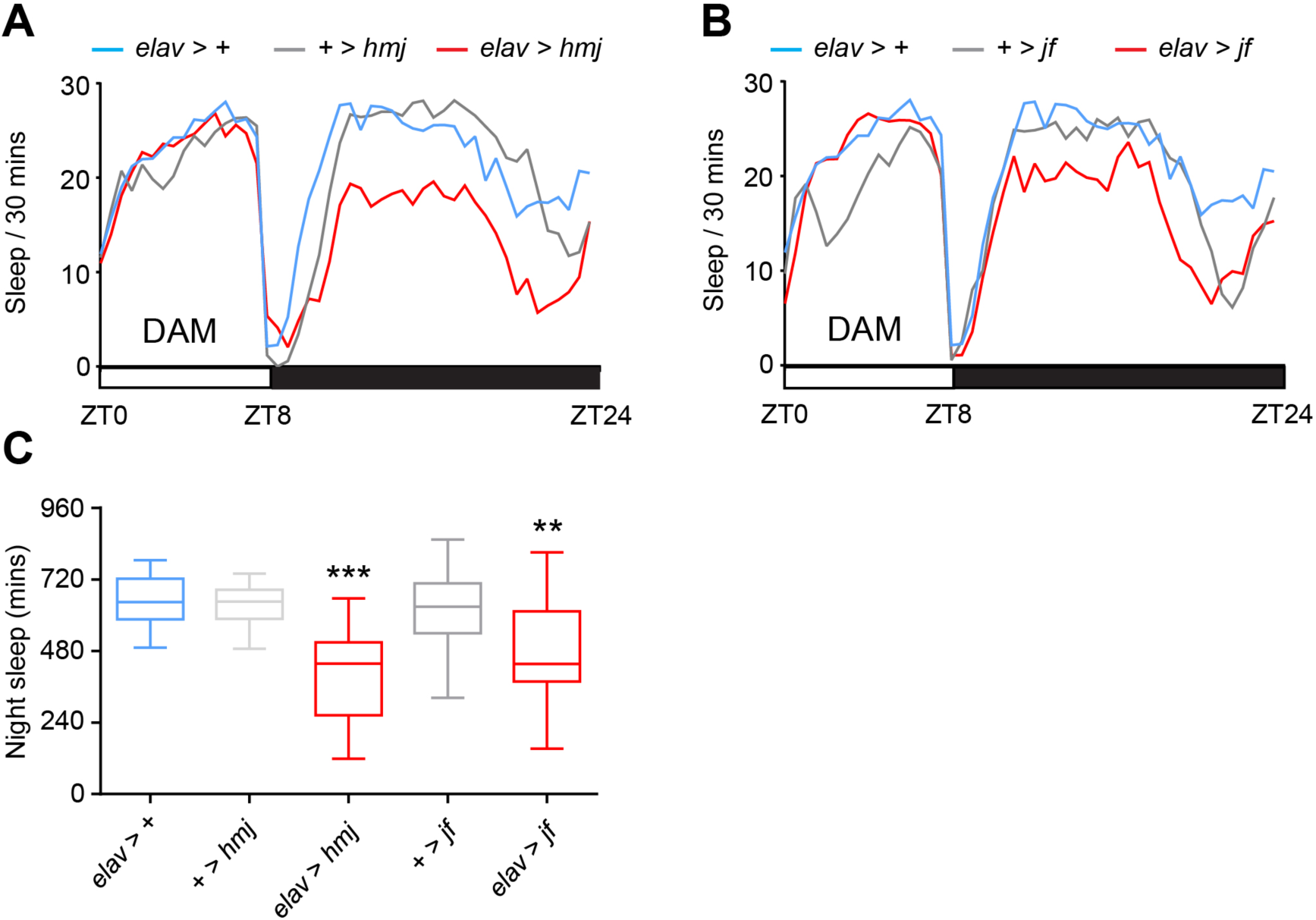
Robust night sleep loss in adult male flies expressing two independent *Nca* RNAi lines *driven by elav*-Gal4. (A-B) Mean sleep profiles under 8L: 16D conditions for *elav*-gal4 drive*, hmj* (A) or *jf* (B) *Nca* RNAi. (C) Median night sleep amounts for genotypes shown in (A-B). *elav > +:* n = 32; *+ > hmj*: n = 26; *elav > hmj*: n = 17; *+ > jf*: n = 32; *elav > jf*: n = 32. **p < 0.01, ***p < 0.001 as compared to driver and RNAi alone controls, Kruskal-Wallis test with Dunn’s post-hoc test.

**Figure 2 supplemental figure 3.**
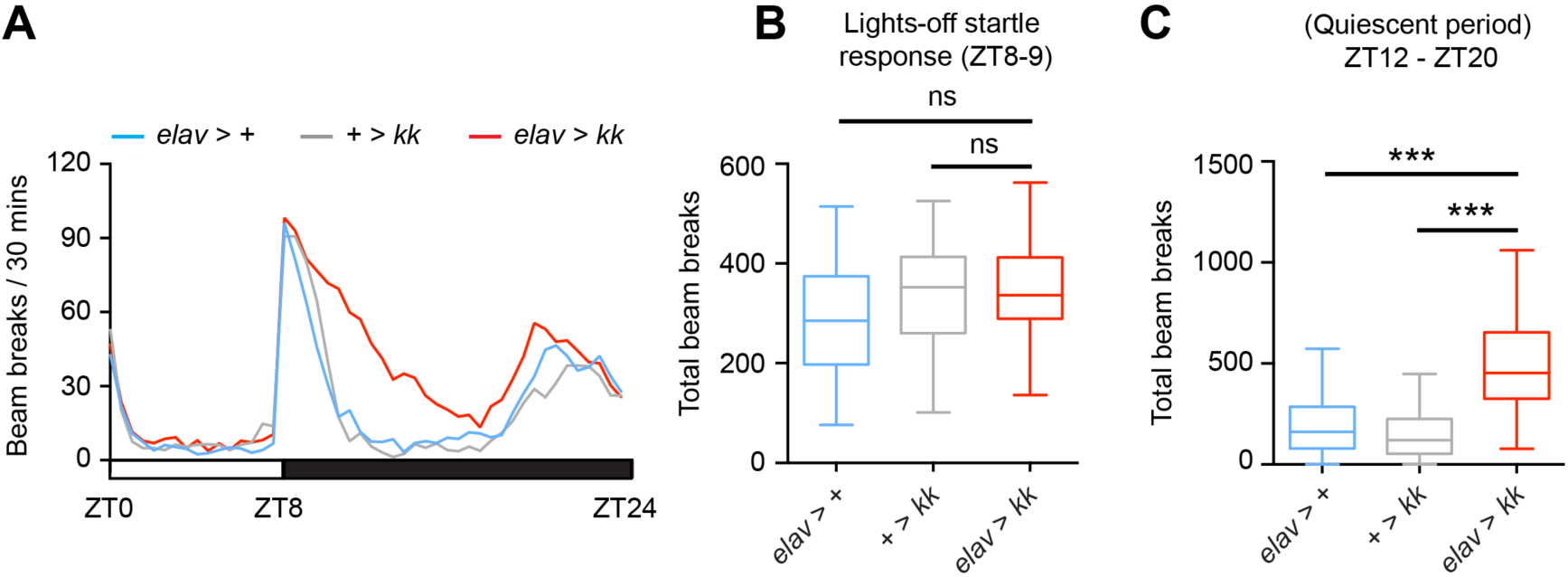
Elevated night time locomotor activity in *Nca*^KD^ flies under 8L: 16D conditions, but no alteration in peak locomotor activity. (A) Mean level of locomotor activity (beam crosses) for *Nca*^KD^ males (*elav > kk*) and associated controls. (B) Median beam breaks during the hour immediately following lights-off (ZT8-9) in the above genotypes. (C) Median total beam crosses during the normally quiescent period of the night (ZT12-20) in the above genotypes. n = 54-55 per genotype. ***p < 0.001, ns – p > 0.05, Mann-Whitney U-test.

**Figure 2 supplemental figure 4.**
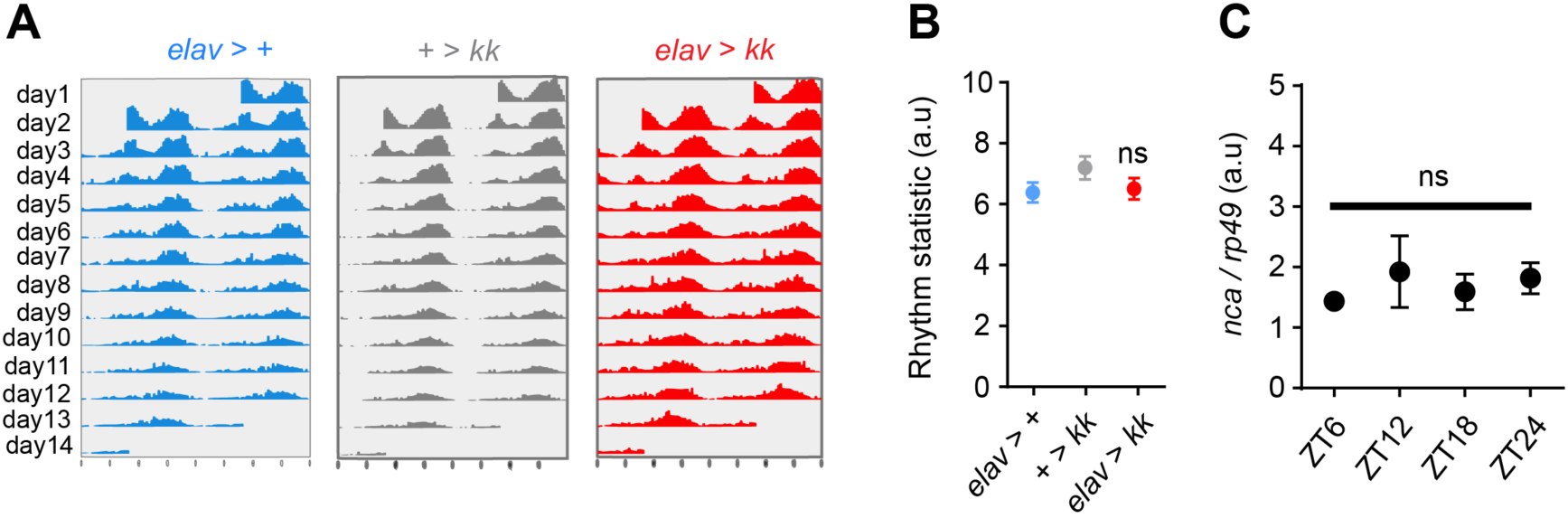
*Nca* knockdown does not alter circadian rhythmicity. (A) Actograms showing representative individual patterns of locomotor activity across two weeks under constant dark conditions. (B) Quantification of locomotor rhythm strength. Robust circadian patterns of locomotor activity were still observed following in adult males expressing *Nca* RNAi (*kk*) under *elav*-Gal4 relative to controls. 15 > n > 14. (C) qPCR analysis of wild type (*Canton-S*) head *Nca* expression across a 24 h period. No evidence of circadian oscillations in *Nca* expression was observed. n = 3 (triplicated qPCR reactions for each time point). Expression levels were normalised to the *rp49* control transcript. ns – p > 0.05, Kruskal-Wallis test with Dunn’s post-hoc test.

**Figure 2 supplemental figure 5.**
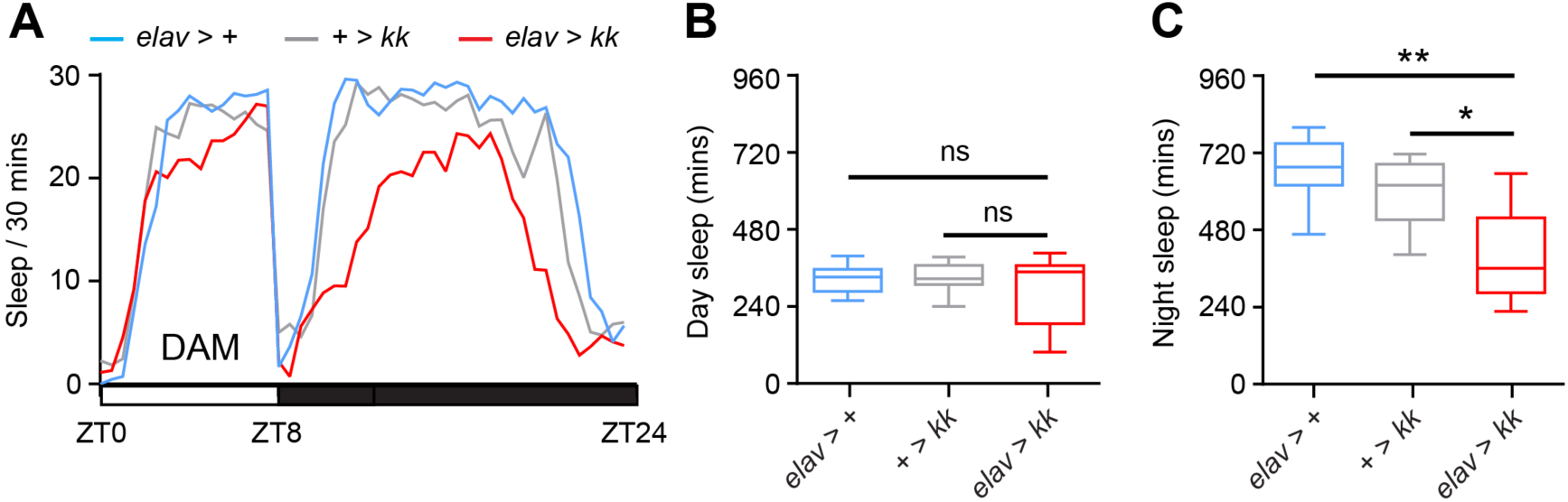
Neuronal *Nca* knockdown results in reduced night sleep in adult virgin female *Drosophila*. (A) Mean sleep patterns of neuronal *Nca* knockdown females (*elav > kk*) and associated controls under 8L:16D conditions. (B-C) Median day sleep is unaffected relative to controls (B), whereas median night sleep is significantly reduced (C). n = 16 per genotype. *p < 0.05, **p < 0.01, Kruskal-Wallis test with Dunn’s post-hoc test.

**Figure 2 supplemental figure 6.**
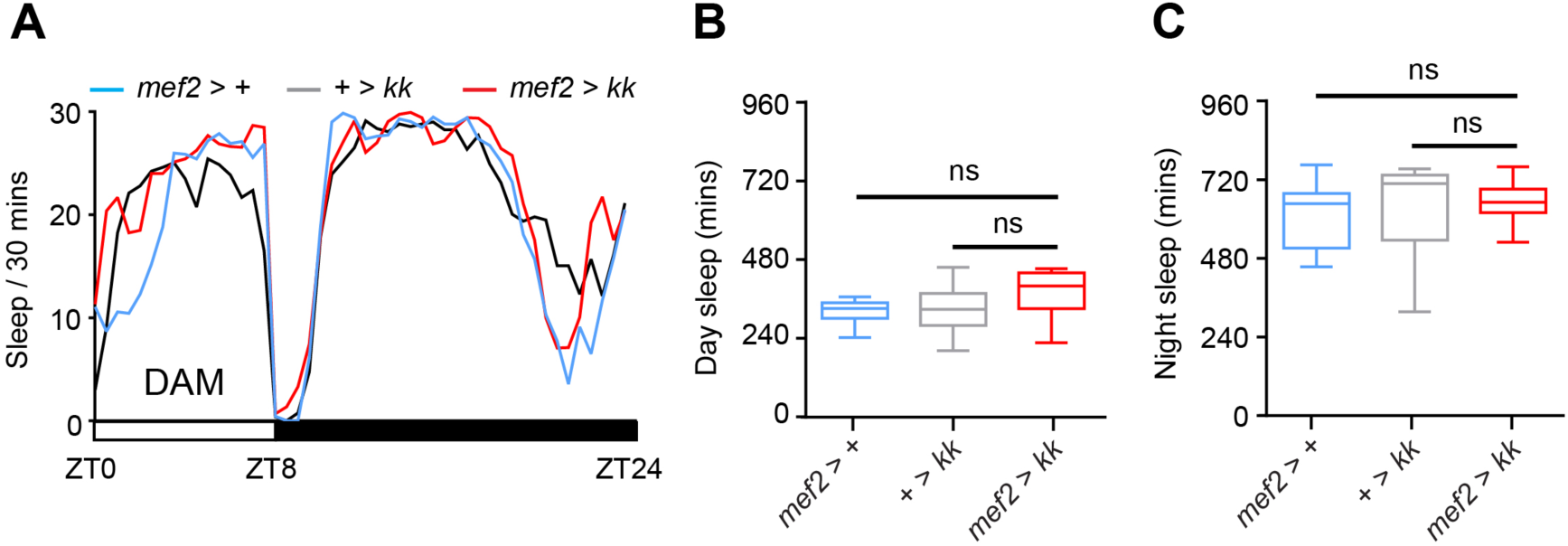
*Nca* knockdown in muscle cells does not affect sleep in *Drosophila*. (A) Mean sleep patterns of adult male flies with muscle-specific *Nca* knockdown via *mef2*-Gal4 (*mef2 > kk*) and associated controls under 8L: 16D. (B-C) Median day (B) and night (C) sleep levels are unaffected relative to controls. n = 16 per genotype. ns – p > 0.05, Kruskal-Wallis test with Dunn’s post-hoc test.

**Figure 2 supplemental figure 7.**
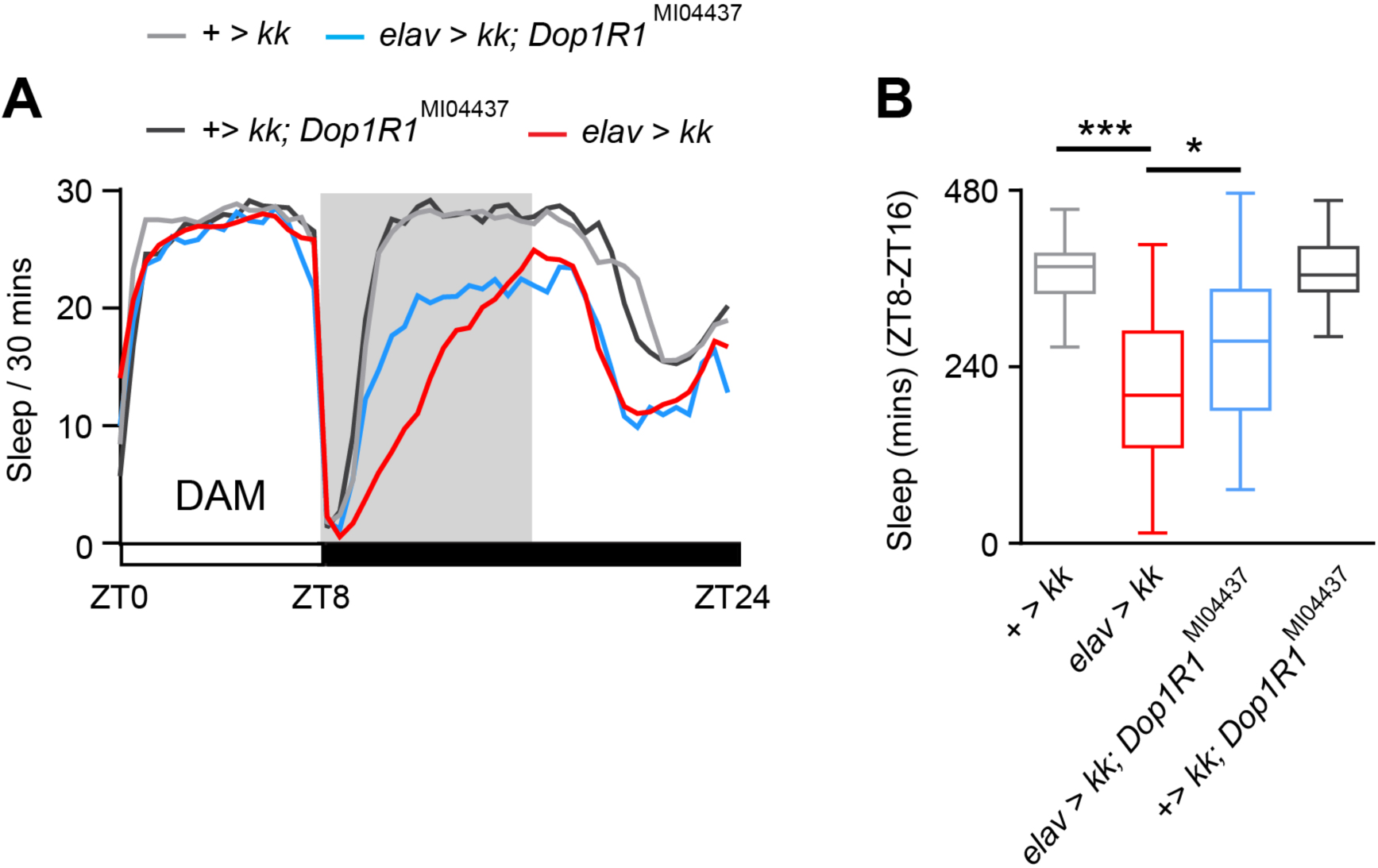
Genetic interaction between *Nca* and a second independent *Dop1R1 allele*. (A) Mean sleep patterns of *Nca*^KD^ males (*elav* > *kk*) and a heterozygous *kk* alone control with and without one copy of the *Dop1R1*^MI04437^ allele under 8L: 16D conditions. *Dop1R1*^MI04437^ is a hypomorphic allele of Dop1R1 that is homozygous viable, and is therefore weaker compared to the homozygous lethal *Dop1R1*^MI03085-GFST.2^ insertion. (B) Median night sleep levels during ZT8-16 are shown. Heterozygosity for *Dop1R1*^MI04437^ had no impact on sleep in *kk* heterozygote controls but partially rescued night sleep during ZT8-16 in *Nca*^KD^ males. *elav* > *kk*: n = 64; + > *kk*: n = 64; *elav* > *kk*; *Dop1R1*^MI04437^/+: n = 39; + > *kk*; *Dop1R1*^MI04437^/+: n = 45. *p < 0.05, ***p < 0.001, Kruskal-Wallis test with Dunn’s post-hoc test.

**Figure 2 supplemental figure 8.**
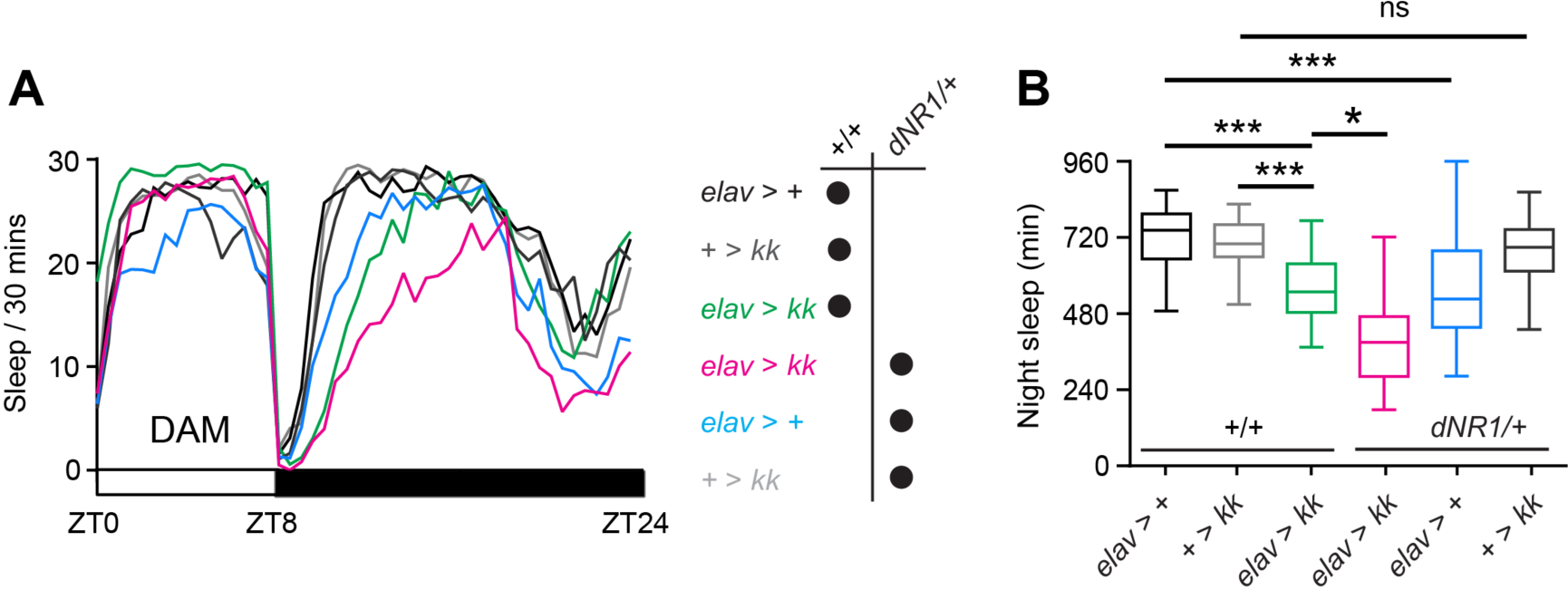
NCA regulate sleeps in a distinct pathway to the dNR1 NMDA receptor. (A) Mean sleep patterns of *Nca*^KD^ males (*elav* > *kk*) and heterozygous controls with and without one copy of the *dNR1*^MI11796^ (*dNR1*/+) allele in 8L: 16D conditions are shown. (B) Median night sleep levels in the above genotypes. Heterozygosity for *dNR1*^MI11796^ resulted in sleep loss in *elav*-Gal4 driver controls, and reduced sleep further in an additive manner in *elav* > *kk Nca*^KD^ males, suggesting that NCA and dNR1 act in separate pathways to promote night sleep. *Elav > kk*; *dNR1*/+: n = 32; *elav > +*; *dNR1*/+: n = 26; *+ > kk*; *dNR1*/+: n = 38; *elav > kk*: n = 32; *elav > +*: n = 30; *+ > kk*: n = 34. ns – p > 0.05, *p < 0.05, **p < 0.01, ***p < 0.001, Kruskal-Wallis test with Dunn’s post-hoc test.

**Figure 3 figure supplement 1.**
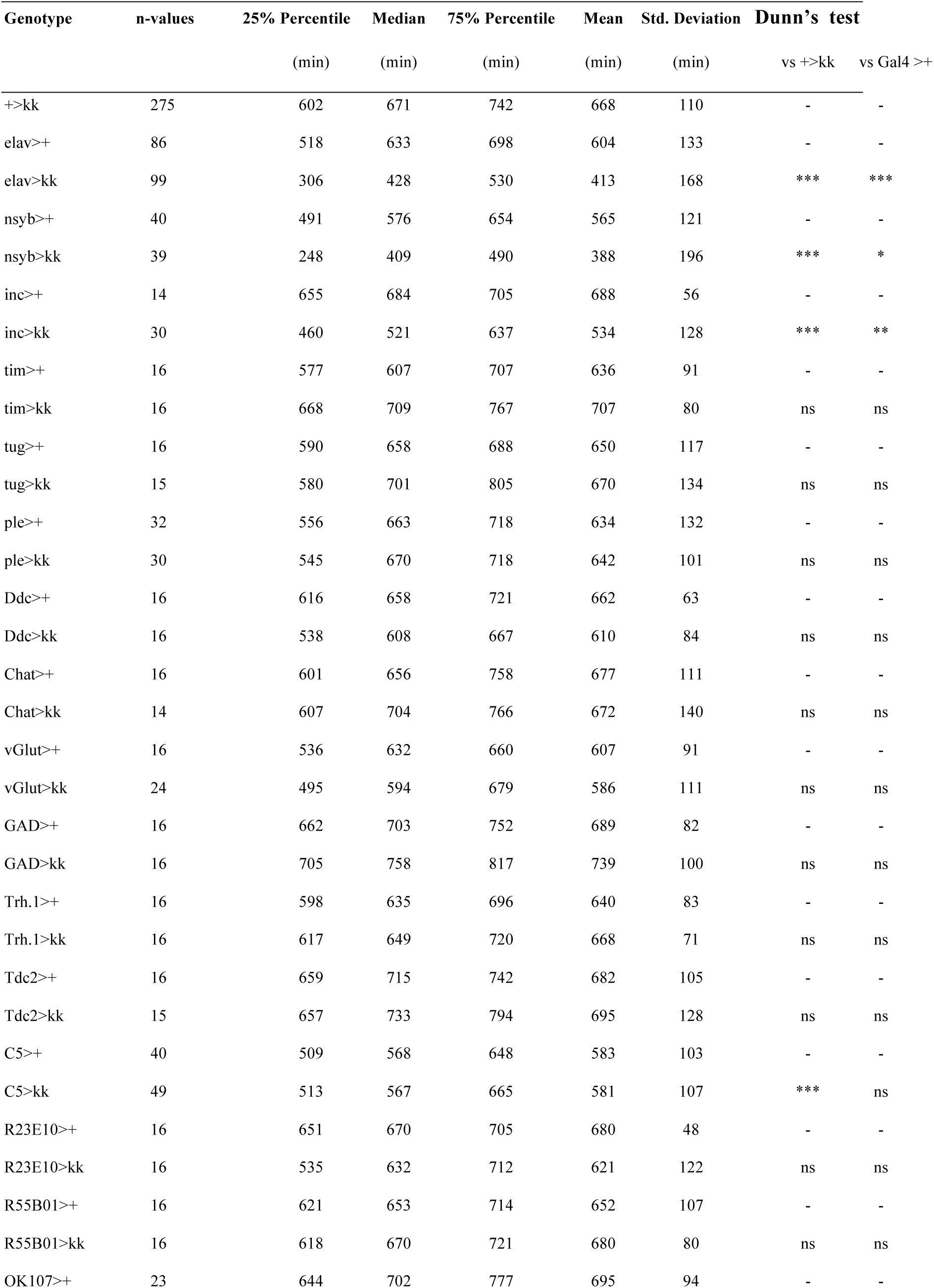
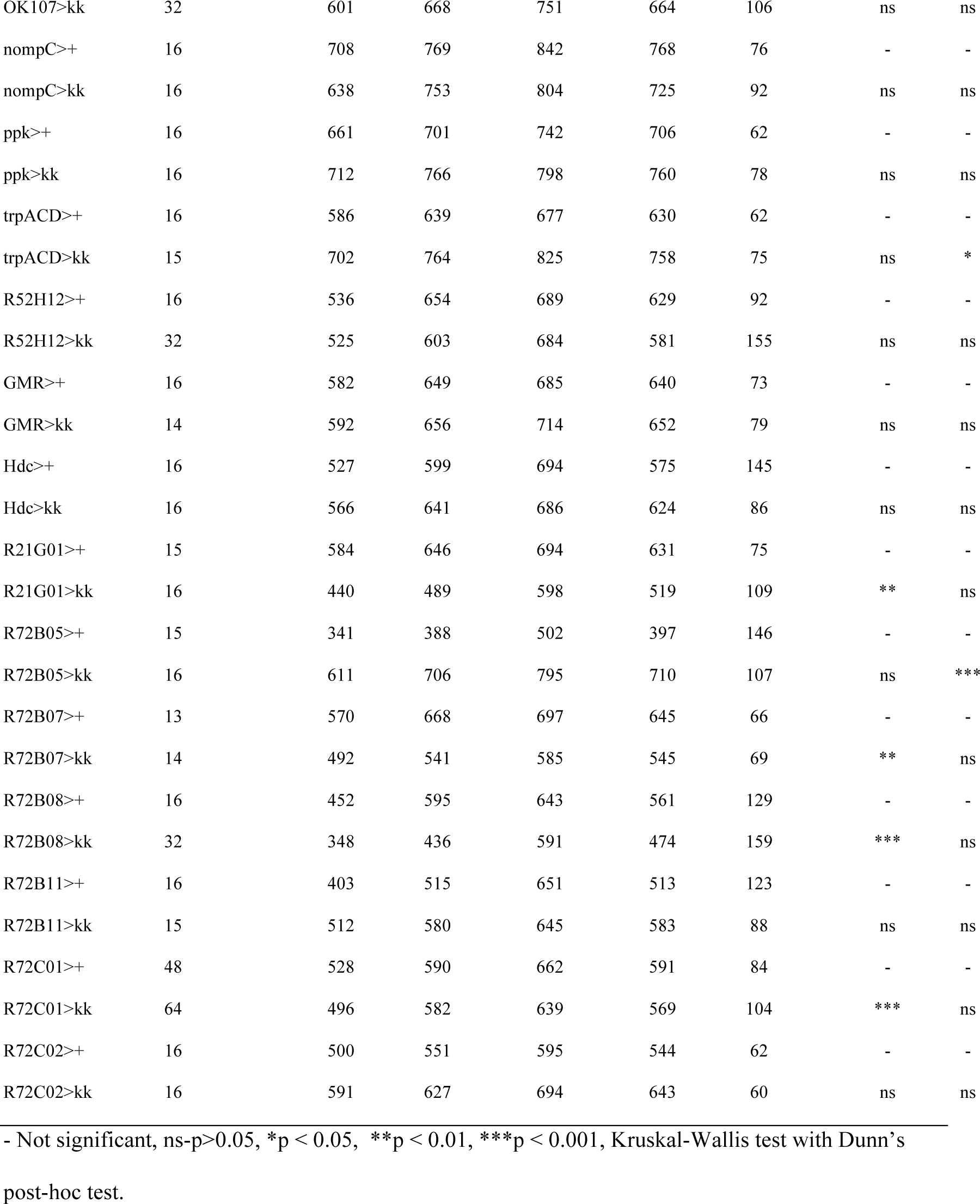
Night sleep levels in flies with *kk Nca RNAi* driven by neuron sub-type-specific Gal4 drivers.

**Figure 3 supplemental figure 2.**
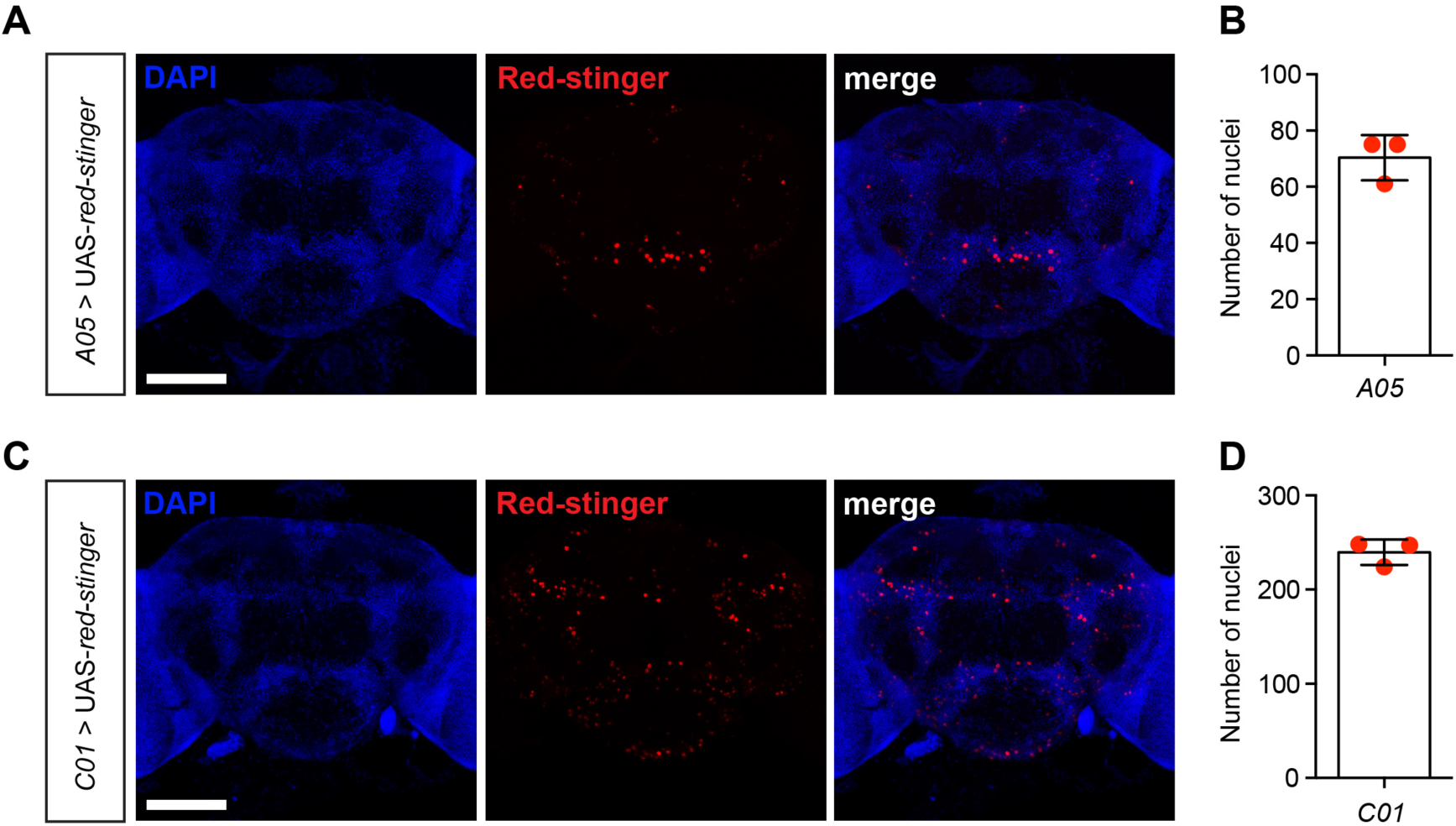
*C01*-Gal4 and *A05*-Gal4 label small subsets of neurons in the adult fly brain. (A, C) Representative confocal z-stacks of adult male brains expressing nuclear RFP marker (Red-stinger) under *A05*-Gal4 (A) or *C01*-Gal4 driver (C). DAPI-staining labels nuclei within the *Drosophila* brain. (B, D) Number of cells expressing Red-stinger driven by *A05-* (B) or *C01*-Gal4 (D). n = 3 for each genotype. Scale bar =100 μm.

**Figure 3 supplemental figure 3.**
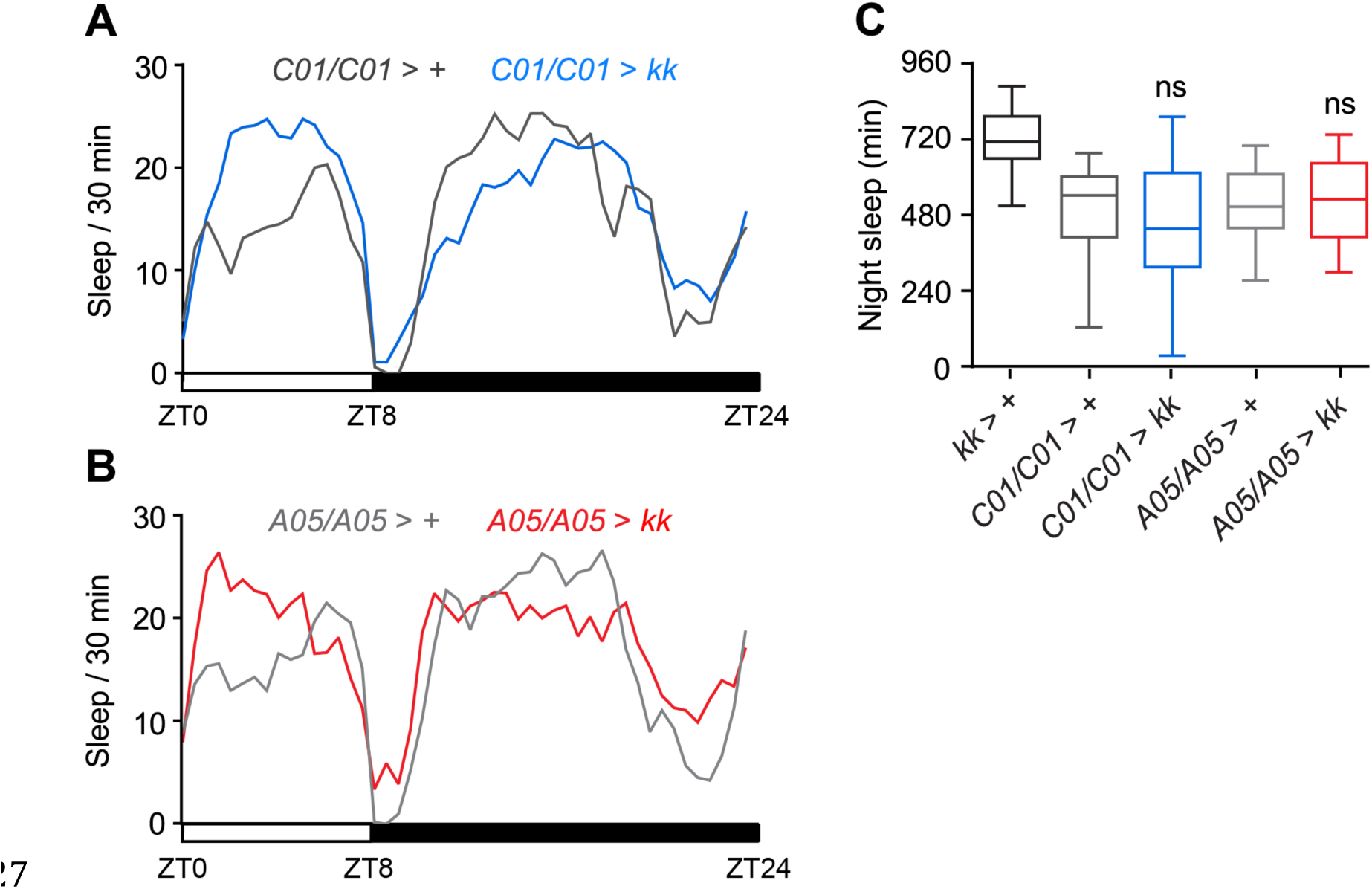
(A) Mean sleep patterns of adult males homozygous for the *C01*-Gal4 driver with and without the *kk Nca* RNAi insertion. (B) Mean sleep patterns of adult males homozygous for the *A05*-Gal4 driver with and without the *kk Nca* RNAi insertion. (C) Median night sleep levels for heterozygous RNAi transgene and homozygous driver controls, and males expressing *Nca* RNAi with two Gal4 driver copies. No night sleep loss was observed using two copies of either driver relative to controls. + > *kk*: n = 80 (the same population was used in Figure 3C, as the experiments were performed side by side), *C01/C01* > +: n = 24, *C01/C01* > *kk*: n = 39; *A05/A05* >: n = 22, *A05/A05* > *kk*: n = 23. ns – p > 0.05, Kruskal-Wallis test with Dunn’s post-hoc test.

**Figure 3 supplemental figure 4.**
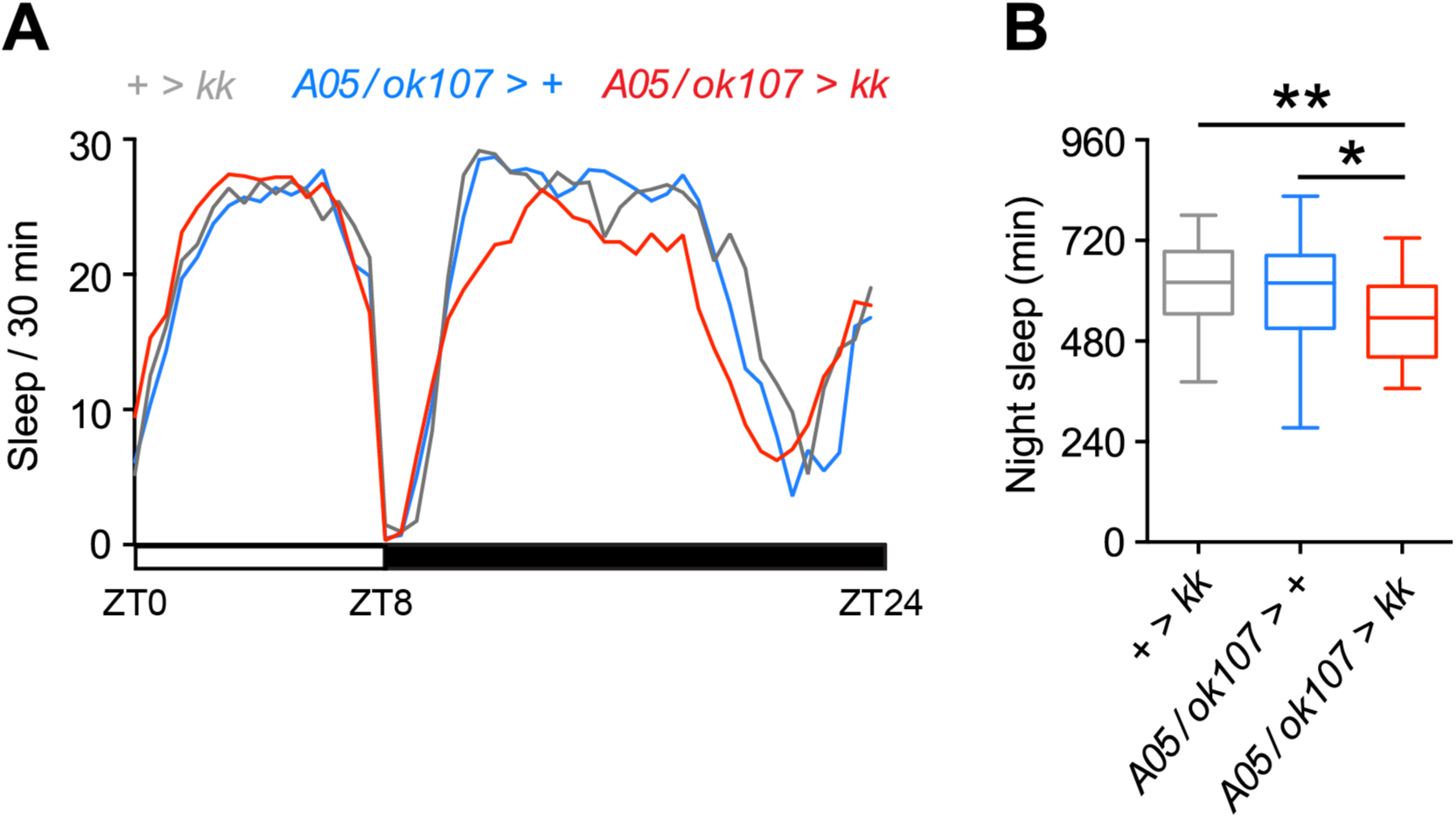
Knockdown of *Nca* in mushroom body and *A05*- neurons results in partial night sleep loss. (A-B) *Nca* knockdown in both *A05* and MB-neurons (defined by *ok107*-Gal4; *ok107*) results in reduced night sleep (also see Figure 3A showing *ok107* > *kk* alone does not cause night sleep loss). Mean sleep patterns in 8L: 16D conditions are shown in (A). Median night sleep levels are shown in (B). + > *kk*: n = 31; *A05/ok107* > +: n = 33; *A05/ok107 > kk*: n = 42. *p < 0.05, **p < 0.01, Kruskal-Wallis test with Dunn’s post-hoc test.

## References

Afonso, D.J., Liu, D., Machado, D.R., Pan, H., Jepson, J.E., Rogulja, D., and Koh, K. (2015). TARANIS Functions with Cyclin A and Cdk1 in a Novel Arousal Center to Control Sleep in *Drosophila*. Curr Biol 25, 1717–1726.

Andrade, R., Foehring, R.C., and Tzingounis, A.V. (2012). The calcium-activated slow AHP: cutting through the Gordian knot. Front Cell Neurosci 6, 47.

Baena-Lopez, L.A., Alexandre, C., Mitchell, A., Pasakarnis, L., and Vincent, J.P. (2013). Accelerated homologous recombination and subsequent genome modification in *Drosophila*. Development 140, 4818–4825.

Braunewell, K.H., and Klein-Szanto, A.J. (2009). Visinin-like proteins (VSNLs): interaction partners and emerging functions in signal transduction of a subfamily of neuronal Ca2+ -sensor proteins. Cell Tissue Res 335, 301–316.

Breakefield, X.O., Blood, A.J., Li, Y., Hallett, M., Hanson, P.I., and Standaert, D.G. (2008). The pathophysiological basis of dystonias. Nat Rev Neurosci 9, 222–234.

Burgoyne, R.D., and Haynes, L.P. (2012). Understanding the physiological roles of the neuronal calcium sensor proteins. Mol Brain 5, 2.

Calabresi, P., Pisani, A., Rothwell, J., Ghiglieri, V., Obeso, J.A., and Picconi, B. (2016). Hyperkinetic disorders and loss of synaptic downscaling. Nat Neurosci 19, 868–875.

Charlesworth, G., Angelova, P.R., Bartolome-Robledo, F., Ryten, M., Trabzuni, D., Stamelou, M., Abramov, A.Y., Bhatia, K.P., and Wood, N.W. (2015). Mutations in HPCA cause autosomal-recessive primary isolated dystonia. Am J Hum Genet 96, 657–665.

Charlesworth, G., Plagnol, V., Holmstrom, K.M., Bras, J., Sheerin, U.M., Preza, E., Rubio-Agusti, I., Ryten, M., Schneider, S.A., Stamelou, M., et al. (2012). Mutations in ANO3 cause dominant craniocervical dystonia: ion channel implicated in pathogenesis. Am J Hum Genet 91, 1041–1050.

Donlea, J.M., Thimgan, M.S., Suzuki, Y., Gottschalk, L., and Shaw, P.J. (2011). Inducing sleepby remote control facilitates memory consolidation in *Drosophila*. Science 332, 1571–1576.

Fahn, S. (1988). Concept and classification of dystonia. Adv Neurol 50, 1–8.

Faville, R., Kottler, B., Goodhill, G.J., Shaw, P.J., and van Swinderen, B. (2015).How deeply does your mutant sleep? Probing arousal to better understand sleep defects in *Drosophila*. Sci Rep 5, 8454.

Fuchs, T., Gavarini, S., Saunders-Pullman, R., Raymond, D., Ehrlich, M.E.,Bressman, S.B., and Ozelius, L.J. (2009). Mutations in the THAP1 gene are responsible for DYT6 primary torsion dystonia. Nat Genet 41, 286–288.

Fuchs, T., Saunders-Pullman, R., Masuho, I., Luciano, M.S., Raymond, D., Factor, S., Lang, A.E., Liang, T.W., Trosch, R.M., White, S., et al. (2013). Mutations in GNAL cause primarytorsion dystonia. Nat Genet 45, 88–92.

Gerfen, C.R., and Surmeier, D.J. (2011). Modulation of striatal projection systems by dopamine. Annu Rev Neurosci 34, 441–466.

Hamada, F.N., Rosenzweig, M., Kang, K., Pulver, S.R., Ghezzi, A., Jegla, T.J., and Garrity, P.A. (2008). An internal thermal sensor controlling temperature preference in *Drosophila*. Nature 454, 217–220.

Helassa, N., Antonyuk, S.V., Lian, L.Y., Haynes, L.P., and Burgoyne, R.D. (2017). Biophysical and functional characterization of hippocalcin mutants responsible for human dystonia. Hum Mol Genet 26: 2426–2435.

Jenett, A., Rubin, G.M., Ngo, T.T., Shepherd, D., Murphy, C., Dionne, H., Pfeiffer,B.D., Cavallaro, A., Hall, D., Jeter, J., et al. (2012). A GAL4-driver line resource for *Drosophila* neurobiology. Cell Rep 2, 991–1001.

Jiang, Y., Pitmon, E., Berry, J., Wolf, F.W., McKenzie, Z., and Lebestky, T.J. (2016). A Genetic Screen To Assess Dopamine Receptor (DopR1) Dependent Sleep Regulation in *Drosophila*. G3 (Bethesda) 6, 4217–4226.

Jo, J., Son, G.H., Winters, B.L., Kim, M.J., Whitcomb, D.J., Dickinson, B.A., Lee, Y.B., Futai, K., Amici, M., Sheng, M., et al. (2010). Muscarinic receptors induce LTD of NMDAR EPSCs via a mechanism involving hippocalcin, AP2 and PSD-95. Nat Neurosci 13, 1216–1224.

Joiner, W.J., Crocker, A., White, B.H., and Sehgal, A. (2006). Sleep in *Drosophila* is regulated by adult mushroom bodies. Nature 441, 757–760.

Kamikouchi, A., Inagaki, H.K., Effertz, T., Hendrich, O., Fiala, A., Göpfert, M.C., andIto, K. (2009). The neural basis of *Drosophila* gravity-sensing and hearing. Nature 458, 165–171.

Karimi, M., and Perlmutter, J.S. (2015). The role of dopamine and dopaminergic pathways in dystonia: insights from neuroimaging. Tremor Other Hyperkinet Mov (N Y) 5, 280.

Kume, K., Kume, S., Park, S.K., Hirsh, J., and Jackson, F.R. (2005). Dopamine is a regulator of arousal in the fruit fly. J Neurosci 25, 7377–7384.

Lamaze, A., Ozturk-Colak, A., Fischer, R., Peschel, N., Koh, K., and Jepson, J.E. (2017). Regulation of sleep plasticity by a thermo-sensitive circuit in *Drosophila*. Sci Rep 7, 40304.

Lebestky, T., Chang, J.S., Dankert, H., Zelnik, L., Kim, Y.C., Han, K.A., Wolf, F.W., Perona, P., and Anderson, D.J. (2009). Two different forms of arousal in *Drosophila* are oppositely regulated by the dopamine D1 receptor ortholog DopR via distinct neural circuits. Neuron 64, 522–536.

Lee, H.J., Weitz, A.J., Bernal-Casas, D., Duffy, B.A., Choy, M., Kravitz, A.V., Kreitzer, A.C., and Lee, J.H. (2016). Activation of Direct and Indirect Pathway Medium Spiny Neurons Drives Distinct Brain-wide Responses. Neuron 91, 412–424.

Lehner, B. (2013). Genotype to phenotype: lessons from model organisms for human genetics. Nat Rev Genet 14, 168–178.

Levine, J.D., Funes, P., Dowse, H.B., and Hall, J.C. (2002a). Advanced analysis of a cryptochrome mutation's effects on the robustness and phase of molecular cycles in isolated peripheral tissues of *Drosophila*. BMC neuroscience 3, 5.

Levine, J.D., Funes, P., Dowse, H.B., and Hall, J.C. (2002b). Signal analysis of behavioral and molecular cycles. BMC neuroscience 3, 1.

Li, Q., Kellner, D.A., Hatch, H.A.M., Yumita, T., Sanchez, S., Machold, R.P., Frank, C. A., and Stavropoulos, N. (2017). Conserved properties of *Drosophila insomniac* link sleep regulation and synaptic function. PLoS Genet 13, e 1006815.

Liu, Q., Liu, S., Kodama, L., Driscoll, M.R., and Wu, M.N. (2012). Two dopaminergic neurons signal to the dorsal fan-shaped body to promote wakefulness in *Drosophila*. Curr Biol 22, 2114–2123.

Liu, S., Lamaze, A., Liu, Q., Tabuchi, M., Yang, Y., Fowler, M., Bharadwaj, R., Zhang, J., Bedont, J., Blackshaw, S., et al. (2014). WIDE AWAKE mediates the circadian timing of sleep onset. Neuron 82, 151–166.

McGary, K.L., Park, T.J., Woods, J.O., Cha, H.J., Wallingford, J.B., and Marcotte,E.M. (2010). Systematic discovery of nonobvious human disease models through orthologous phenotypes. Proc Natl Acad Sci U S A 107, 6544–6549.

Mencacci, N.E., Rubio-Agusti, I., Zdebik, A., Asmus, F., Ludtmann, M.H., Ryten,M., Plagnol, V., Hauser, A.K., Bandres-Ciga, S., Bettencourt, C., et al. (2015). A missense mutation in KCTD17 causes autosomal dominant myoclonus-dystonia. Am J Hum Genet 96, 938–947.

Miesenbock, G. (2012). Synapto-pHluorins: genetically encoded reporters of synaptic transmission. Cold Spring Harb Protoc 2012, 213–217.

Nitabach, M.N., Blau, J., and Holmes, T.C. (2002). Electrical silencing of *Drosophila* pacemaker neurons stops the free-running circadian clock. Cell 109, 485–495.

Pappas, S.S., Darr, K., Holley, S.M., Cepeda, C., Mabrouk, O.S., Wong, J.M., LeWitt, T.M., Paudel, R., Houlden, H., Kennedy, R.T., et al. (2015). Forebrain deletion of the dystonia protein torsinA causes dystonic-like movements and loss of striatal cholinergic neurons. Elife 4, e08352.

Park, D., and Griffith, L.C. (2006). Electrophysiological and anatomical characterization of PDF-positive clock neurons in the intact adult *Drosophila* brain. J Neurophysiol 95, 3955–3960.

Peterson, D.A., Sejnowski, T.J., and Poizner, H. (2010). Convergent evidence for abnormal striatal synaptic plasticity in dystonia. Neurobiol Dis 37, 558–573.

Pfeiffenberger, C., and Allada, R. (2012). Cul3 and the BTB adaptor insomniac are key regulators of sleep homeostasis and a dopamine arousal pathway in *Drosophila.* PLoS Genet 8, e1003003.

Pfeiffenberger, C., Lear, B.C., Keegan, K.P., and Allada, R. (2010a). Locomotor activity level monitoring using the *Drosophila* Activity Monitoring (DAM) System. Cold Spring Harb Protoc 2010, pdb prot5518.

Pfeiffenberger, C., Lear, B.C., Keegan, K.P., and Allada, R. (2010b). Processing sleep data created with the *Drosophila* Activity Monitoring (DAM) System. Cold Spring Harb Protoc 2010, pdb prot5520.

Pitman, J.L., McGill, J.J., Keegan, K.P., and Allada, R. (2006). A dynamic role for the mushroom bodies in promoting sleep in *Drosophila*. Nature 441, 753–756.

Rogulja, D., and Young, M.W. (2012). Control of sleep by cyclin A and its regulator. Science 335, 1617–1621.

Seidner, G., Robinson, J.E., Wu, M., Worden, K., Masek, P., Roberts, S.W., Keene, A.C., and Joiner, W.J. (2015). Identification of Neurons with a Privileged Role in Sleep Homeostasis in *Drosophila melanogaster*. Curr Biol 25, 2928–2938.

Shi, M., Yue, Z., Kuryatov, A., Lindstrom, J.M., and Sehgal, A. (2014). Identification of Redeye, a new sleep-regulating protein whose expression is modulated by sleep amount. Elife 3, e01473.

Sitaraman, D., Aso, Y., Jin, X., Chen, N., Felix, M., Rubin, G.M., and Nitabach, M.N. (2015). Propagation of Homeostatic Sleep Signals by Segregated Synaptic Microcircuits of the *Drosophila* Mushroom Body. Curr Biol 25, 2915–2927.

Stavropoulos, N., and Young, M.W. (2011). insomniac and Cullin-3 regulate sleep and wakefulness in Drosophila. Neuron 72, 964–976.

Tanabe, L.M., Kim, C.E., Alagem, N., and Dauer, W.T. (2009). Primary dystonia: molecules and mechanisms. Nat Rev Neurol 5, 598–609.

Tecuapetla, F., Jin, X., Lima, S.Q., and Costa, R.M. (2016). Complementary Contributions of Striatal Projection Pathways to Action Initiation and Execution. Cell 166, 703–715.

Teng, D.H., Chen, C.K., and Hurley, J.B. (1994). A highly conserved homologue of bovine neurocalcin in *Drosophila melanogaster* is a Ca(2+)-binding protein expressed in neuronal tissues. J Biol Chem 269, 31900–31907.

Tomita, J., Ban, G., and Kume, K. (2017). Genes and neural circuits for sleep of the fruit fly. Neurosci Res 118, 82–91.

Tomita, J., Ueno, T., Mitsuyoshi, M., Kume, S., and Kume, K. (2015). The NMDA Receptor Promotes Sleep in the Fruit Fly, *Drosophila melanogaster*. PLoS One 10, e0128101.

Tzingounis, A.V., Kobayashi, M., Takamatsu, K., and Nicoll, R.A. (2007). Hippocalcin gates the calcium activation of the slow afterhyperpolarization in hippocampal pyramidal cells. Neuron 53, 487–493.

Ueno, K., Naganos, S., Hirano, Y., Horiuchi, J., and Saitoe, M. (2013). Long-term enhancement of synaptic transmission between antennal lobe and mushroom body in cultured *Drosophila*brain. J Physiol 591, 287–302.

Ueno, T., Tomita, J., Tanimoto, H., Endo, K., Ito, K., Kume, S., and Kume, K. (2012). Identification of a dopamine pathway that regulates sleep and arousal in *Drosophila*. Nat Neurosci 15, 1516–1523.

Wakabayashi-Ito, N., Ajjuri, R.R., Henderson, B.W., Doherty, O.M., Breakefield, X.O., O'Donnell, J.M., and Ito, N. (2015). Mutant human torsinA, responsible for early-onset dystonia, dominantly suppresses GTPCH expression, dopamine levels and locomotion in *Drosophila melanogaster*. Biol Open 4, 585–595.

Wakabayashi-Ito, N., Doherty, O.M., Moriyama, H., Breakefield, X.O., Gusella, J.F., O'Donnell, J.M., and Ito, N. (2011). Dtorsin, the *Drosophila* ortholog of the early-onset dystonia TOR1A (DYT1), plays a novel role in dopamine metabolism. PLoS One 6, e26183.

Weisheit, C.E., and Dauer, W.T. (2015). A novel conditional knock-in approach defines molecular and circuit effects of the DYT1 dystonia mutation. Hum Mol Genet 24, 6459–6472.

Wenning, G.K., Kiechl, S., Seppi, K., Muller, J., Hogl, B., Saletu, M., Rungger, G., Gasperi, A., Willeit, J., and Poewe, W. (2005). Prevalence of movement disorders in men and women aged 50-89 years (Bruneck Study cohort): a population-based study. Lancet Neurol 4, 815–820.

Wu, M., Robinson, J.E., and Joiner, W.J. (2014). SLEEPLESS is a bifunctional regulator of excitability and cholinergic synaptic transmission. Curr Biol 24, 621–629.

Yokoi, F., Dang, M.T., Liu, J., Gandre, J.R., Kwon, K., Yuen, R., and Li, Y. (2015). Decreased dopamine receptor 1 activity and impaired motor-skill transfer in Dyt1 DeltaGAG heterozygous knock-in mice. Behav Brain Res 279, 202–210.

Zhong, L., Bellemer, A., Yan, H., Ken, H., Jessica, R., Hwang, R.Y., Pitt, G.S., and Tracey, W.D. (2012). Thermosensory and nonthermosensory isoforms of *Drosophila melanogaster* TRPA1 reveal heat-sensor domains of a thermoTRP Channel. Cell Rep 1, 43–55.

